# The multiscale architecture of trait aggression: from meta-analytic convergence to multimodal characterization

**DOI:** 10.64898/2026.09.26.753333

**Authors:** Adili Abudukadier, Shounuo Chen, Keying Jiang, Fanxi Ding, Quanyu Kan, Liyuan Gu, Pinchun Wang, Mingxing Cheng, Haili Zhang, Shizhen Yan, Qiang Wang

## Abstract

**Background:** Trait aggression shows substantial interindividual variability, yet its neurobiological architecture across structural, functional, molecular, and macroscale organizational levels remains poorly understood.

**Methods:** We developed a convergent multiscale framework to characterize the neurobiological architecture associated with trait aggression. We first integrated a large-sample voxel-based morphometry analysis (n = 922) with published neuroimaging studies using two complementary coordinate-based meta-analytic approaches, activation likelihood estimation and seed-based d mapping (total N = 3,671). Convergent meta-analytic findings were subsequently mapped onto functional connectivity networks, and their molecular signatures were characterized using imaging transcriptomics and neurotransmitter mapping. Finally, we examined whether the spatial organization of aggression-related networks was embedded within intrinsic multimodal brain hierarchies spanning morphometric, functional, transcriptomic, and neurochemical dimensions.

**Results:** Across complementary analyses, trait aggression converged on a reproducible structural–functional network centered on the dorsomedial prefrontal cortex and dorsal anterior cingulate cortex. This architecture exhibited distinct molecular signatures, including gene-expression patterns enriched for synaptic functions and robust spatial associations with serotonergic, dopaminergic, and noradrenergic transporter systems. Moreover, the spatial organization of the aggression-related functional network was systematically predicted by the brain’s intrinsic multimodal gradients, indicating that it is embedded within broader principles of cortical hierarchical organization.

**Conclusions:** These findings move beyond the localization of aggression to discrete brain regions and identify a reproducible multiscale architecture linking aggression-related structural variation to functional network organization, molecular signatures, and the brain’s intrinsic multimodal hierarchy. This integrative framework provides a systems-level account of the neurobiological basis of individual differences in trait aggression.

## Introduction

Aggressive behavior is a common social behavior that, in evolutionary terms, has adaptive value by protecting resources and supporting survival(Feltes & De Boer, 2021; Nelson & Trainor, 2007). However, when aggression exceeds social norms and becomes frequent, intense, and driven by the intent to harm, it poses a significant public health challenge. At the individual level, aggression is not just a momentary reaction but a stable, trait-like attribute(Allen & Anderson, 2017; Granecki, 2014). Trait aggression reflects an individual’s consistent, cross-situational tendencies in cognition (hostility), emotion (anger), and behavior (physical and verbal aggression; Buss & Perry, 1992; Buss & Warren, 2000), And it represents the core psychological construct for understanding individual differences in aggression. Elucidating the neurobiological basis of trait aggression is therefore essential for clarifying the vulnerability mechanisms of aggression, identifying high-risk individuals, and developing effective intervention strategies.

Existing studies have preliminarily delineated gray matter volume (GMV) abnormality patterns associated with aggression. Key correlates include the medial prefrontal cortex (mPFC; Coccaro et al., 2018; Dugré & De Brito, 2024), orbitofrontal cortex (Wang et al., 2024), and anterior cingulate cortex (ACC; Hostetler et al., 2024). Complementary evidence from animal models, optogenetics, and human deep brain stimulation further implicates the amygdala(Aubry et al., 2025; Roberts et al., 2021; Rosell & Siever, 2015), hypothalamus, striatum, and prefrontal cortex(Aubry et al., 2025; Feltes & De Boer, 2021; Roberts et al., 2021; Rosell & Siever, 2015; Santin, 2025). Nevertheless, these findings are marked by considerable heterogeneity due to variations in sample size (n < 500), analytical methods, comorbidities, and population characteristics. Consequently, consistent structural markers of trait aggression in healthy populations still lack systematic validation. Moreover, structural localization alone cannot fully explain how aggression translates from brain to behavior. Gray matter alterations likely exert distal effects through functional connectivity networks(Boes et al., 2015; Peng et al., 2022). Prior work has begun to characterize these networks from several angles: GMV abnormality networks (thalamic–dorsal and salience networks; Chen et al., 2025), task-evoked activation abnormality networks (basal ganglia network; Chen et al., 2025; Wang et al., 2024), and resting-state activity abnormality networks (limbic-prefrontal network; Rosell & Siever, 2015). Yet, a more robust and core structural–functional network mapping remains to be established.

Imaging transcriptomics and multi-receptor association analyses act as critical bridges between molecular features and macroscopic phenotypes. These approaches have seen important advances in other psychiatric disorders(Han et al., 2025; Liu et al., 2025; Wang et al., 2026; Wang et al., 2026; Xia et al., 2022; Yang et al., 2021), but remain in their infancy for aggression research. If aggression-related networks possess biological specificity, their spatial distribution should correlate with distinct gene expression profiles and neurotransmitter receptor distributions(Hansen et al., 2022). Thus, applying imaging transcriptomics and multi-receptor association analyses to these networks can anchor macroscopic features to microscopic molecular mechanisms, offering deeper constraints on the biological basis of aggression.

Behavioral tendencies are shaped by the synergistic influence of multiscale and multimodal brain features. Converging evidence indicates that the cerebral cortex is organized along continuous gradients that progress from primary sensorimotor regions to transmodal association cortices(Huntenburg et al., 2018; Margulies et al., 2016). This hierarchical organization is evident not only in functional connectivity and morphology but also in multimodal gradients of gene expression and neurotransmitter receptor distributions(Burt et al., 2018). However, it remains unknown how the organizational pattern of aggression-related networks aligns with this macroscale hierarchical architecture.

Fig. 1 illustrates our methodological framework. We propose that trait aggression is linked to a specific structural–functional network, which exhibits a distinct molecular signature (gene expression and receptor profiles) and is hierarchically constrained by the brain’s multimodal architecture. First, voxel-based morphometry (VBM) analysis on a large sample (n = 922) identified aggression-related GMV changes, which were integrated into an existing VBM meta-analytic framework. Subsequently, meta-analyses using activation likelihood estimation (ALE) and seed-based *d* mapping (SDM) yielded consistent cross-study structural correlates. Peak coordinates of the significant clusters from these meta-analyses were then used as seeds for functional connectivity network mapping (FCNM), generating ALE FCNM and SDM FCNM maps and thus translating structural findings into functional networks. Imaging transcriptomics and multi-receptor association analyses were applied to the four resulting z maps (the ALE map, the SDM map, the ALE FCNM map, and the SDM FCNM map) to extract underlying gene expression and neurotransmitter receptor profiles. In parallel, we constructed four whole-brain modality gradients—a morphometric similarity gradient (MSG), a functional connectivity gradient (FCG), a transcriptomic similarity gradient (TSG), and a receptor similarity gradient (RSG) —to represent the brain’s intrinsic multimodal architecture. Separately, we derived aggression-specific network gradients from the ALE FCNM and SDM FCNM maps. Finally, multiple linear regression was performed with the four whole-brain gradients as predictors and the aggression-specific network gradients as the dependent variable, aiming to reveal the hierarchical biological factors shaping the cortical organization of the aggression related functional network. Collectively, this integrates individual level structural findings, meta analytic consensus, network mapping, molecular signatures, and multimodal gradients into a unified framework for understanding the multimodal brain mechanisms of aggression.

**Fig. 1.**
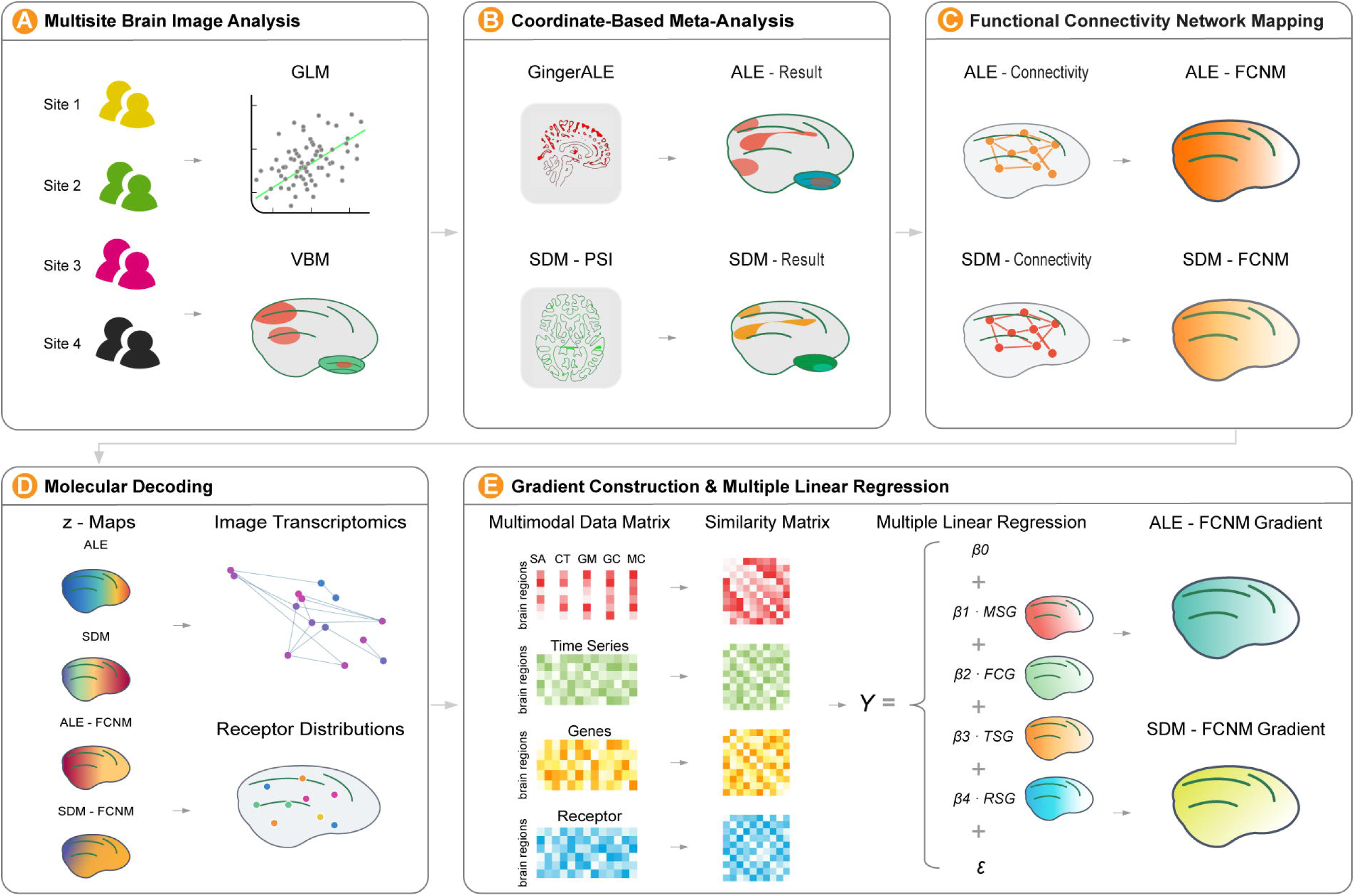
Schematic overview of the analytic workflow. (A) Multisite Brain Image Analysis; (B) Coordinate-Based Meta-Analysis; (C) Functional Connectivity Network Mapping; (D) Molecular Decoding, including imaging transcriptomics and receptor association analysis; (E) Gradient Construction and Multiple Linear Regression

## Methods

### Participants and Data preparation

Participants were recruited from universities via bulletin boards. A total of 922 college students had complete T1-weighted imaging data and Buss–Warren Aggression Questionnaire (BWAQ, 34 items, Cronbach’s α = 0.91; responses (*M*_age_ = 19.98 years, *SD* = 1.67; 561 females; Buss & Warren, 2000), and of these, 753 also provided resting-state fMRI data (442 females). This study was approved by the Ethics Review Committee.

Detailed data acquisition and preprocessing parameters are provided in Appendix 1.

### VBM Analysis

GMV was analyzed voxel-wise using the optimized FSL-VBM pipeline(Jenkinson et al., 2012; Smith et al., 2004) (http://fsl.fmrib.ox.ac.uk/fsl/fslwiki/FSLVBM). A whole-brain general linear model was constructed with trait aggression score as the variable of interest, and sex, age, paternal education level, and total intracranial volume (TIV) were included as covariates. Statistical inference was carried out using the *flameo* command with 5,000 permutations and threshold-free cluster enhancement (TFCE); significant voxels were identified at a cluster-level corrected threshold of *p* < 0.01.

### Meta-Analysis

Following common meta-analytic methods(Cheung & Vijayakumar, 2016), under the PRISMA framework (Fig. 5), we systematically searched for and included 12 VBM studies on trait aggression in healthy populations. Together with the VBM results from the present study, 13 studies (3671 individuals, Table 1) were submitted to coordinate-based meta-analyses using both activation likelihood estimation (ALE) and seed-based *d* mapping (SDM; for details on the search strategy and methodology, see Appendix 2).

**Table 1.** Characteristics of included studies in the meta-analysis.

| Study | Space | Software | N | Correction | Measurement |
| --- | --- | --- | --- | --- | --- |
| (Besteher et al., 2017) | MNI | SPM12 | 409 | FDR < 0.05 | SCL-90-R |
| (Breitschuh et al., 2018) | MNI | SPM12 | N<sub>1</sub>=26<br>N<sub>2</sub>=21 | uncorrected $p < 0.001$ | FAF |
| (Chester et al., 2017) | MNI | FSL | 138 | $p < 0.05$, FWE-corrected (TFCE) | BAQ |
| (Coccaro et al., 2018) | MNI | SPM8 | 287 | $p < 0.05$, FWE-corrected | LHA |
| (Gong et al., 2023) | MNI | SPM8 | 176 | $p < 0.05$, FWE-corrected (TFCE) | MS & SAS |
| (Gong et al., 2024) | MNI | SPM12 | 185 | $p < 0.05$, FWE-corrected (TFCE) | BPAQ |
| (Gorka et al., 2015) | MNI | SPM8 | 464 | $p < 0.05$, FWE-corrected | BPAQ |
| (Raschle et al., 2018) | MNI | SPM12 | 189 | $p < 0.05$, FWE-corrected (TFCE) | ICU & YPI |
| (Seidenbecher et al., 2023) | MNI | SPM12 | 57 | $p < 0.05$, FWE-corrected (TFCE) | BPAQ |
| (Wang et al., 2023) | MNI | SPM12 | 202 | $p < 0.05$, FWE-corrected (TFCE) | BPAQ |
| (Wei & Xia, 2023) | MNI | SPM12 | 325 | $p < 0.05$, FWE-corrected (TFCE) | SEIP-Q |
| (Zhu et al., 2019) | MNI | SPM8 | 240 | $p < 0.05$, FWE-corrected (TFCE) | RPQ |
| The current study | MNI | FSL | 922 | $p < 0.01$, FWE-corrected (TFCE) | BWAQ |

Note. SCL-90-R, Aggression subscale of The Symptom Checklist-90 (Derogatis et al., 1973);
FAF: Factors of Aggression Questionnaire (Hampel, 1975); RPQ: Reactive-proactive aggression
questionnaire (Dodge & Coie, 1987; Raine et al., 2006); BAQ, Brief Aggression Questionnaire
(Webster et al., 2015); LHA, Life History of Aggression (Coccaro et al., 1997); MS,
Machiavellianism Subscale; SAS, Social Aggression Subscale; ICU, Inventory of
Callous-Unemotional traits (Essau et al., 2006); YPI, Youth Psychopathic traits Inventory
(Andershed et al., 2007); BPAQ, Buss-Perry Aggression Questionnaire (Buss & Perry, 1992);
SEIP-Q, Social Emotional Information Processing questionnaire (Coccaro et al., 2017); BWAQ,
Buss-Warren Aggression Questionnaire (Buss & Warren, 2000).

#### ALE meta-analysis

The latest revised algorithm of GingerALE (3.0.2) was used to model 3D probability distributions of the reported foci, quantifying topographic convergence across studies(Eickhoff et al., 2009; Turkeltaub et al., 2002). The resulting ALE scores were compared against an empirical null distribution derived from 1,000 permutations, and cluster-level familywise error (FWE) correction was applied with TFCE (*p* < 0.05). The robustness of the results was evaluated through jackknife analysis.

#### SDM-PSI meta-analysis

Seed-based *d* mapping with permutation of subject images (SDM-PSI v6.23) was employed to synthesize both peak coordinates and their effect sizes(Albajes-Eizagirre et al., 2019). Significance was determined using an uncorrected voxel-level threshold of *p_uncorrected_* < 0.001 with a cluster extent of ≥ 10 voxels(Dugré et al., 2020). Residual heterogeneity was assessed with the *I²* statistic, and publication bias was examined with Egger’s test. Jackknife sensitivity analysis was also conducted to confirm the stability of the findings.

### Functional Connectivity Network Mapping (FCNM)

Using the 4 cluster peak coordinates from the ALE meta-analysis and the 3 cluster peak coordinates from the SDM meta-analysis as seeds, we constructed seed-to-whole-brain functional connectivity maps based on our preprocessed resting-state fMRI data with AFNI 23.0.07. For each seed, the correlation coefficient map was *Fisher-z* transformed; the resulting *z*-maps were then averaged separately across the ALE seeds and the SDM seeds to generate the ALE-FCNM and SDM-FCNM whole-brain functional connectivity maps. Each map was subsequently entered into a voxel-wise one-sample general linear model, and significance was determined using *randomise* (5,000 permutations, voxel-based FWE correction, *p* < 0.05). Robustness analyses are reported in Appendix 3.

### Imaging Transcriptomics Analysis

Microarray gene expression data were obtained from the AHBA and processed following the standard pipeline, including mapping to a 300-region cortical parcellation(Schaefer et al., 2018), probe re-annotation, and cross-donor normalization. Partial least squares regression (PLSR) was used to separately relate the spatial variability of the four maps (ALE, SDM, ALE-FCNM, SDM-FCNM) to gene expression profiles. The significance of PLSR components was assessed with 5,000 permutations (*p_perm_* < 0.05). The association between significant components and trait aggression was tested using 5,000 spin permutations (*p_spin_* < 0.05). Gene weights were determined for significance through 1,000 bootstrap iterations (*p_Bonferroni_* < 0.05).

### Receptor Distribution Association Analysis

Using the toolbox implementing the approach of Hansen et al(Hansen et al., 2022), the whole-brain three-dimensional normative atlas of 19 receptors and transporters was mapped to the 300-region cortical parcellation. For each receptor, Pearson correlations were computed between its spatial distribution and each of the four imaging maps (*p_Bonferroni_* < 0.05).

### Multimodal Gradient Construction and Association Analysis

To extract multimodal cortical organization gradients, we constructed six 300 × 300 inter-regional similarity matrices, all based on the Schaefer 300 region parcellation.

#### Morphometric similarity matrix

Based on cortical reconstruction from the T1-weighted data of this study, five morphometric features (SA, surface area; CT, cortical thickness; GM, gray matter; GC, Gaussian curvature; MC, mean curvature) were extracted to build an inter-regional morphometric similarity matrix.

#### Functional connectivity matrix

The group-averaged functional connectivity matrix from the Human Connectome Project (HCP), provided in the Brainspace toolbox(Vos de Wael et al., 2020), was used.

#### ALE-FCNM and SDM-FCNM network matrices

For each of the two FCNM maps, symmetric Kullback–Leibler divergence was computed between every pair of regions to generate a 300-region average connectivity matrix.

#### Gene expression and receptor distribution matrices

Inter-regional similarity matrices were separately calculated from the preprocessed gene expression profiles and the cortical distributions of the receptors.

For each matrix, the top 10% of connections per node were retained, and cosine similarity between nodes was subsequently computed. Diffusion embedding was performed using the Brainspace toolbox to extract gradients for each modality. These gradients were then aligned across modalities through iterative Joint Embedding (10 iterations) with a random reference, yielding aligned principal gradients for each modality.

To quantify the predictive contributions of multimodal gradients to the organization of aggression-related functional networks, we used the MSG, FCG (from HCP), TSG, and RSG as predictors, with the ALE-FCNM principal gradient and the SDM-FCNM principal gradient serving as dependent variables in separate multiple linear regression models (*p_Bonferroni_*< 0.05). Parameter estimation of multiple linear regression was conducted using the ordinary least squares (OLS) method, and HC3 (Heteroscedasticity-consistent standard errors), Bootstrap, Permutation, and outlier robust methods were employed for robustness analysis (Table S6.1).

### Sensitivity Analysis

To examine the robustness of the results, the following analyses were performed: (i) In the VBM analysis, sex, age, paternal education, and total intracranial volume (TIV) were included as covariates to verify the stability of the linear regression results. (ii) A comparative analysis was performed using both ALE and SDM methods, with a leave one study out (jackknife) analysis conducted for each approach (see Appendix 2 for details). (iii) For the FCNM, an independent dataset (*n* = 111) was used to validate the reliability of both ALE FCNM and SDM FCNM. (iv) In the imaging transcriptomics analysis, the robustness of the PLSR components was assessed using permutation tests (5,000 permutations). The correlation between PLS1 and the aggression related brain network features was evaluated using spin permutation (5,000 permutations). The gene weights in PLS1 were examined via bootstrap resampling (1,000 iterations). (v) Four comparative imaging transcriptomics and receptor distribution association analyses were performed, each based on different aggression related brain network features: structural networks derived from ALE and from SDM, and functional networks derived from ALE FCNM and from SDM FCNM. (vi) For the aggression related specific network gradient, two analytical pipelines were constructed based on ALE FCNM and SDM FCNM, respectively, and the mutually validated results from the two pipelines were taken as the robust findings of this study. (vii) When building the multiple linear regression model, several common approaches were employed to test the stability, including HC3 standard errors, bootstrap (1,000 iterations), permutation (1,000 iterations), and outlier robust methods.

## Results

### Morphological Substrates of Trait Aggression via VBM Analysis

To verify the morphological substrates underlying trait aggression in a multi-site large sample (n = 922), we performed a voxel-wise analysis of whole-brain GMV while controlling for sex, age, paternal education level, and total intracranial volume (TIV). We found that trait aggression was positively associated with GMVs in the dorsomedial prefrontal cortex (dmPFC; peak MNI x/y/z = −14/44/44, z = 4.53), frontal pole (FP; peak MNI x/y/z = 4/62/18, *z* = 4.58), middle temporal gyrus (MTG; peak MNI x/y/z = 52/−26/−16, *z* = 5.10), lateral occipital cortex (LOC; peak MNI x/y/z = 34/−62/34, *z* = 5.60), and cerebellum (peak MNI x/y/z = −6/−52/−20, *z* = 5.70) (Fig. 2A), suggesting that these regions are involved in trait aggression.

**Fig. 2.**
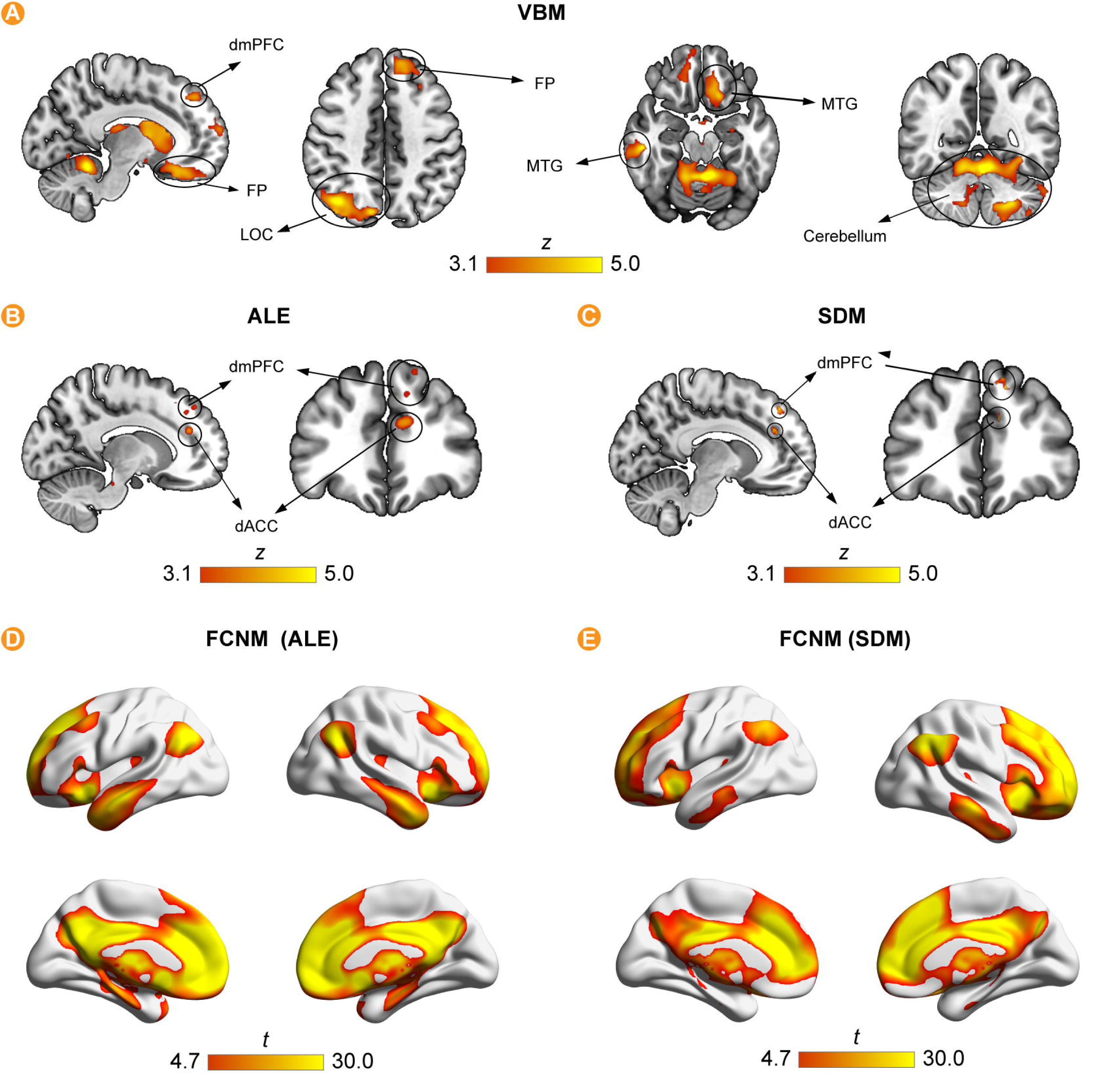
The results of VBM, meta-analysis, and FCNM. (A) VBM z-map, p < 0.01, FWE-corrected (TFCE); (B) ALE z-map, p < 0.05, FWE-corrected (TFCE); (C) SDM z-map, p < 0.001, uncorrected; (D) ALE-FCNM t-map, p < 0.05, FWE-corrected; (E) SDM-FCNM t-map, p < 0.05, FWE-corrected.

### Coordinate-based Meta-analyses of GMV Using ALE and SDM

Although we identified neural correlates of trait aggression in a large sample, it remains unclear whether these structural characteristics are consistent across studies. Given different coordinate-based meta-analytic methods with distinct algorithmic emphases, we employed two mainstream approaches—ALE and SDM—for cross-validation and comparison. Results revealed that significant ALE clusters encompassed the anterior cingulate cortex (ACC), dmPFC, posterior cingulate cortex (PCC), and dorsolateral prefrontal cortex (dlPFC) (Fig. 2B), whereas SDM identified the ACC and dmPFC (Fig. 2C). Notably, conjunction analysis further indicated that the two methods converged in the dmPFC (SDM: peak MNI x/y/z = −14/40/42, *z* = 3.91; ALE: peak MNI x/y/z = −14/44/44, *z* = 3.90) and dorsal ACC (SDM: peak MNI x/y/z = −10/38/28, *z* = 3.68; ALE: peak MNI x/y/z = −10/38/22, *z* = 4.21). Furthermore, the heterogeneity analysis and publication bias of the SDM results indicated that the heterogeneity was low and the publication bias was not significant (see Table S2.1 for details). The jackknife analysis of ALE and SDM results also showed good robustness (see Figure S2.2 and Table S2.2 for details). The details on the literature searching strategy, methodology and results of robustness analysis can be found in Appendix 2.

### Aggression-related Functional Connectivity Network

Here, we used the significant cluster peak coordinates from the ALE and SDM meta-analyses as seeds and performed seed-based FCNM in the discovery dataset (n = 753) and validation dataset (n = 111), yielding the ALE-FCNM (Fig. 2D) and SDM-FCNM (Fig. 2E) results. The regions showing convergent co-connectivity for ALE-FCNM included the cerebellum, left thalamus, caudate, insula, PCC, and dorsal ACC; for SDM-FCNM, these regions included the Frontal Pole, dorsal ACC, dmPFC, PCC, frontal orbital cortex, lateral occipital cortex, cerebellum, insula, caudate, and thalamus (*p_FWE_*< 0.05). Both maps primarily implicated the default mode network (DMN) and the frontoparietal network (FPN) in trait aggression (see Appendix 3 for robustness analyses).

### Transcriptional Mechanisms: Spatial Association of Gene Expression

Next, we explored the potential molecular mechanisms underlying the aggressive brain network. Using left-hemisphere gene expression data (150 × 15,362) from the Allen Human Brain Atlas (AHBA), we performed PLSR to separately relate the spatial variability of the four previously defined statistical maps (ALE, SDM, ALE-FCNM, SDM-FCNM) to gene expression profiles (Significant genes are reported in Appendix 4). Results showed that the first PLS component (PLS1) was significant for ALE (explained variability (*EV)* = 21.60%, *p*_perm_ = 0.003; *r* = −0.40, *p_spin_* < 0.001), ALE-FCNM (*EV* = 28.07%, *p_perm_* = 0.002; *r* = −0.16, *p_spin_* = 0.003), and SDM-FCNM (*EV* = 42.21%, *p_perm_* < 0.001; *r* = −0.28, *p_spin_*< 0.001), but not for SDM (*EV* = 16.14%, *p_perm_* = 0.070; *r* = −0.06, *p_spin_* < 0.001; see Fig. 3AC for the details). Component 1 represented a transcriptional profile characterized by higher expression in the ACC and temporal pole (TP) regions and lower expression in the occipital cortex. Notably, its high expression is associated with lower aggressive traits. For subsequent enrichment analyses, we took the intersection of upregulated genes (significant positive weights, *p*_Bonferroni_ < 0.05) and downregulated genes (significant negative weights, *p_Bonferroni_* < 0.05) from the ALE-related transcriptional results (ALE and ALE-FCNM); for SDM-related results, only the significant (*p_Bonferroni_* < 0.05) upregulated and downregulated genes from SDM-FCNM were used.

**Fig. 3.**
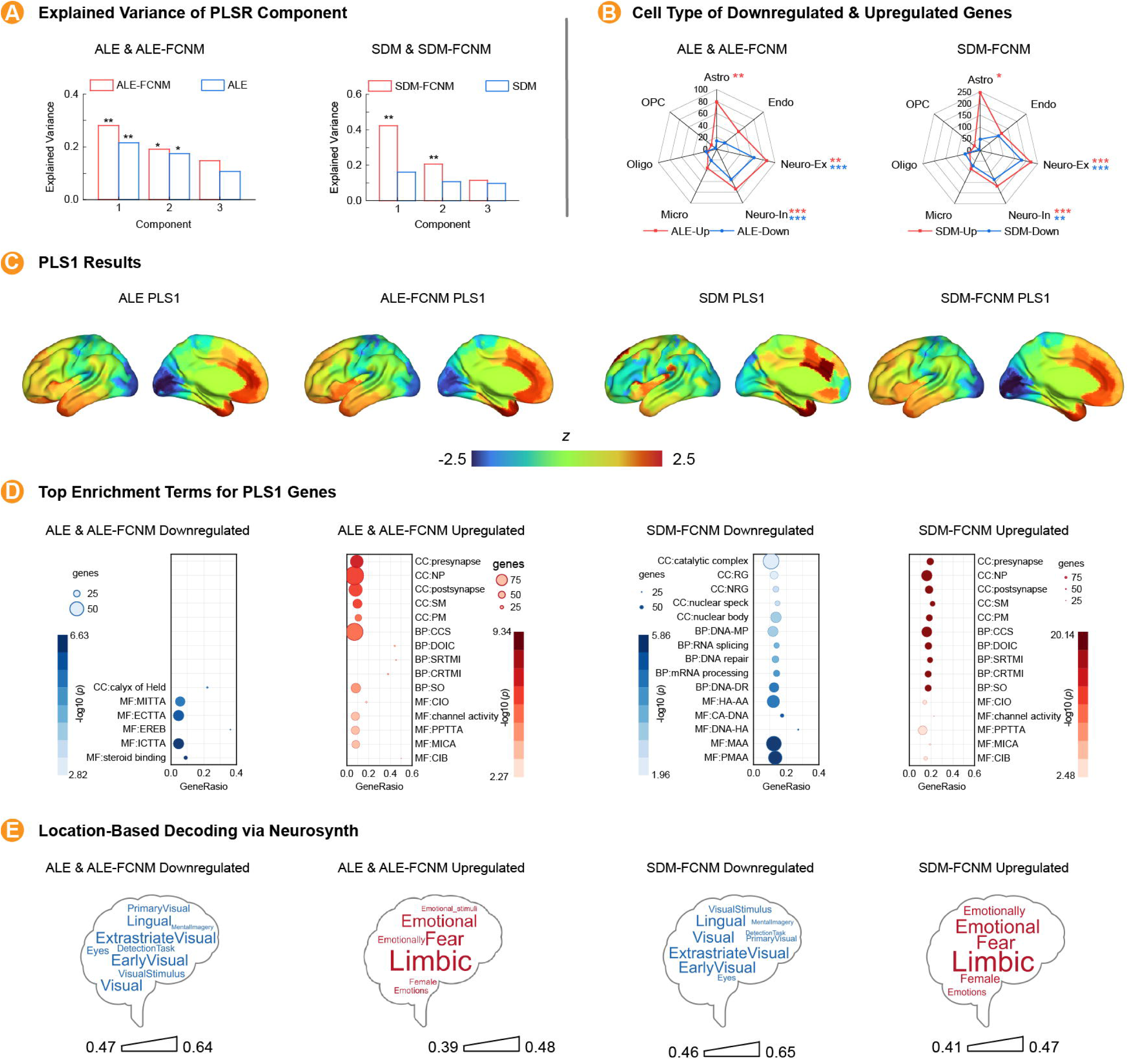
Transcriptomics analysis results. (A) PLSR components and explained variance; (B) Cell types that related to downregulated genes (blue) and upregulated genes (red); (C) Brain region mapping from component 1 of ALE, ALE-FCNM, SDM, SDM-FCNM based PLSR; (D) GO enrichment analysis of the aggression-related genes (top 5 was demonstrated) see Table S4.5 for full GO terms and other details; (E) Location-based Decoding via Neurosynth; *p* < 0.05, *p* < 0.01, *p* < 0.001.

Overall, the upregulated genes converged on synaptic signaling-related pathways, whereas the downregulated genes were more heterogeneous. Specifically, the upregulated genes from the ALE- and SDM-related analyses showed high consistency in molecular function terms such as monoatomic ion channel activity, biological process terms such as cell–cell signaling and synapse organization, and cellular component terms such as presynapse, neuron projection, synaptic membrane, and postsynapse, whereas the downregulated genes were more variable (Fig. 3D). Cell-type analysis revealed that upregulated genes from both the ALE and SDM contexts were significantly associated with inhibitory neurons (Neuro-In), excitatory neurons (Neuro-EX) and astrocytes (Astro), while downregulated genes were associated with inhibitory neurons and excitatory neurons (*p_FDR_* < 0.05, Fig. 3B). Spatiotemporal analysis showed that upregulated genes exhibited more continuous associations with the thalamus in early–mid prenatal periods, the striatum in mid-prenatal periods, and the amygdala and cortex in late prenatal periods; downregulated genes showed more continuous associations with the cerebellum in mid-to-late prenatal periods and with the cortex and thalamus in late prenatal periods. Functional decoding with Neurosynth (Yarkoni et al., 2011) showed that upregulated genes were related to emotion-associated cognitive functions, whereas downregulated genes were related to visual processing (Fig. 3E).

### Receptor Fingerprint: The Association of Neurotransmitter Distribution

Finally, to further characterize brain changes associated with trait aggression, we examined their correspondence with the distribution of specific neurotransmitters (see Fig. 4A). Based on the ALE, ALE FCNM, and SDM FCNM z maps, aggression related regions showed significant positive associations (*p_Bonferroni_* < 0.05) with serotonergic (5 HT1a/2a/4), dopaminergic (D1/DAT), histaminergic (H3), cannabinoid (CB1), opioid (MOR), and glutamatergic (mGluR5) systems, whereas a significant negative correlation was found with the noradrenergic transporter (NET). The results of the tripartite map show that the brain distribution patterns of serotonergic, dopaminergic, and noradrenergic transporter systems are robust across the three modalities of aggressive brain networks (Fig. 4B). For the SDM z map, the associations were not significant (see Appendix 5 for details).

**Fig. 4.**
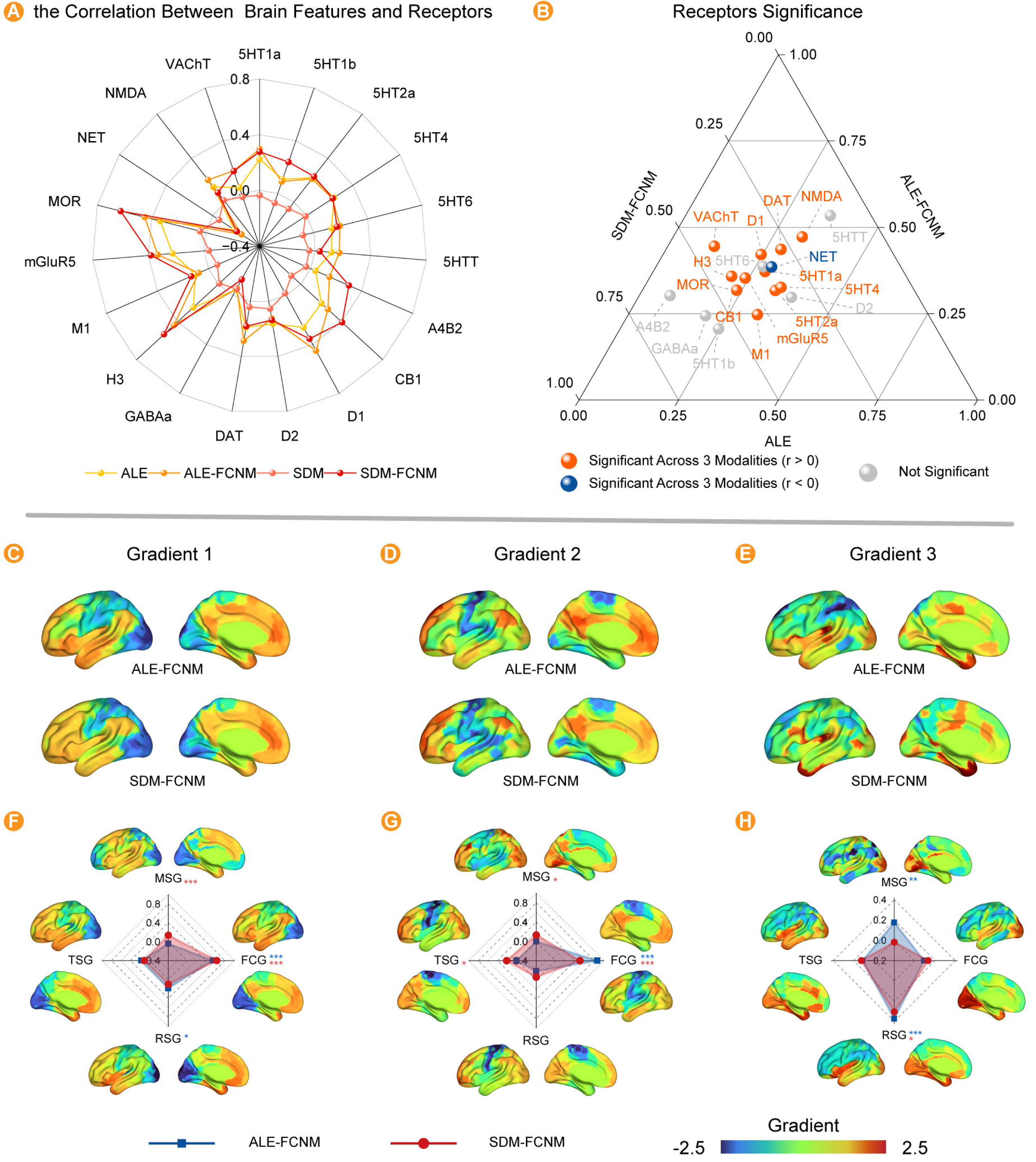
Receptor-association analysis, Gradient construction and multiple linear regression results. (A) the Correlation Between Brain Features and Receptors; (B) Receptors Significance Across 3 Modalities; the position of each receptor is determined by its normalized correlation coefficients across the three methods (sum = 1). Proximity to the triangle center indicates high cross-modal robustness; (C) Gradient 1 of ALE-FCNM (up) and SDM-FCNM (down); (D) Gradient 2 of ALE-FCNM (up) and SDM-FCNM (down); (E) Gradient 3 of ALE-FCNM and SDM-FCNM; (F) Gradient 1 of MSG (up), FCG (right), RCG (down), GCG (left) and the coefficient of multiple linear regression; (G) Gradient 2 of MSG, FCG, RCG, GCG and the coefficient of multiple linear regression; (H) Gradient 3 of MSG, FCG, RSG, GSG and the coefficient of multiple linear regression. Note: *p* < 0.05, *p* < 0.01, *p* < 0.001.

**Fig. 5.**
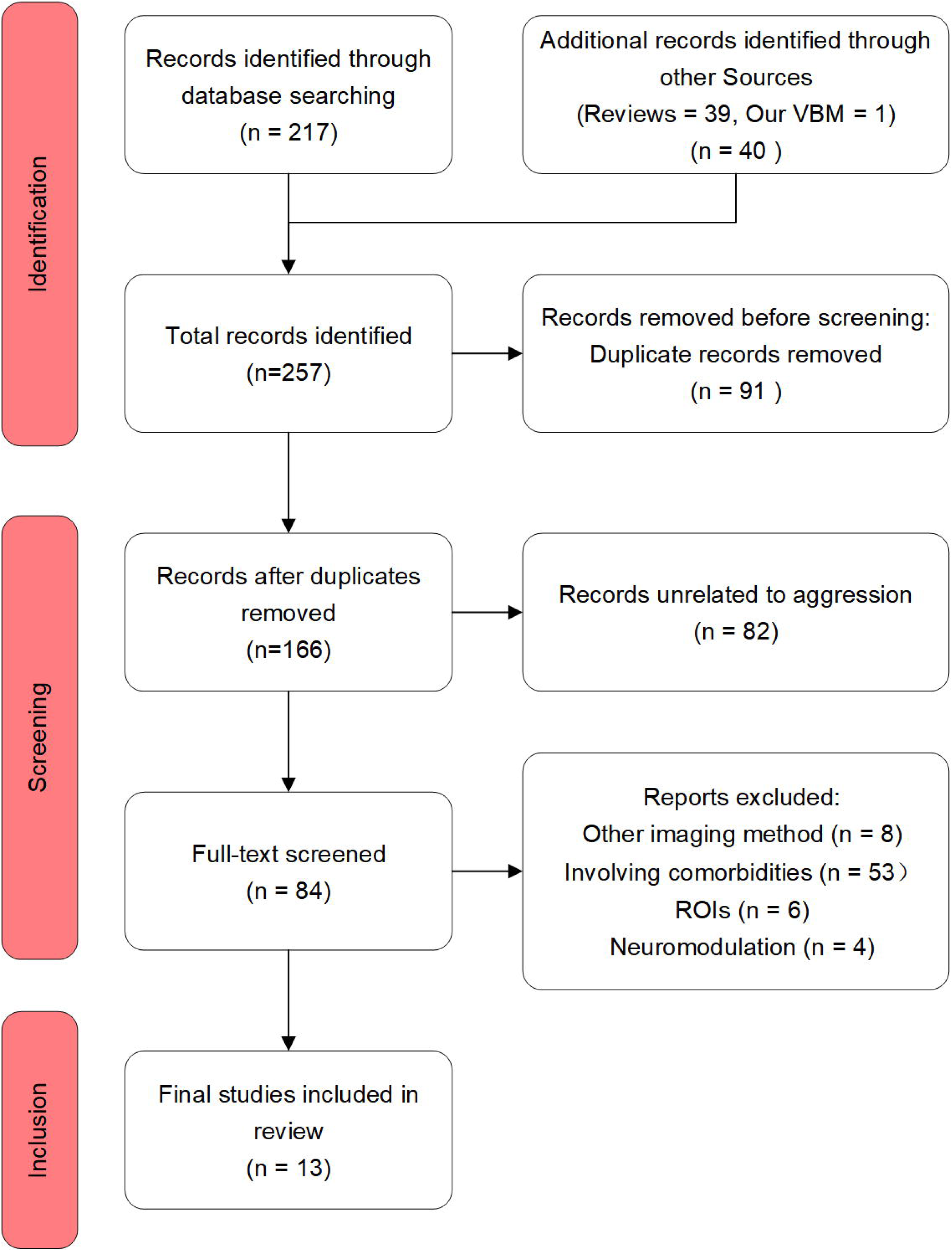
The PRISMA flowchart.

### The Organizational Gradient of the Aggressive Function Network and Multimodal Prediction

To investigate the relationship between the organizational characteristics of aggression-related functional connectivity networks and the brain’s intrinsic hierarchical organization, we used MSG, FCG, TSG, and RSG as predictors to fit aggression-related network gradients (ALE-FCNM gradients and SDM-FCNM gradients) using multiple linear regression. Fig. 4 (the second part) presents the results of the main Gradients and multiple linear regression (gradient1: Fig. 4CF; gradient 2: Fig. 4DG; gradient 3: Fig. 4EH). The results showed that the four gradient patterns significantly predicted the aggression-related network gradients (ALE-FCNM gradients: *F_gradient1_* (4, 295) = 167.455, *p_Bonferroni_* < 0.001, *Adj.R²* = 0.69; *F_gradient2_* (4, 295) = 105.27, *p_Bonferroni_* < 0.001, *Adj.R²* = 0.58; *F_gradient3_* (4, 295) = 50.59, *p_Bonferroni_* < 0.001, *Adj.R²* = 0.40; SDM-FCNM gradients: *F_gradient1_* (4, 295) = 281.49, *p_Bonferroni_* < 0.001, *Adj.R²* = 0.79; *F_gradient2_* (4, 295) = 73.32, *p_Bonferroni_* < 0.001, *Adj.R²* = 0.50; *F_gradient3_* (4, 295) = 26.87, *p_Bonferroni_* < 0.001, *Adj.R²* = 0.26). Among these, FCG was consistently significant in gradients 1 and 2 (for both the ALE-FCNM and SDM-FCNM gradients), while RSG was consistently significant in gradient 3 (see Appendix 6 for robustness analyses). This finding ultimately demonstrates that the cortical distribution of aggression-related functional networks can be effectively predicted by the brain’s multi-level organizational principles, providing a unified graded framework for understanding individual differences in trait aggression.

## Discussion

To characterize the neurobiological architecture of trait aggression, we integrated large-sample structural neuroimaging and meta-analysis with functional network mapping, molecular profiling, and multimodal cortical gradients, providing convergent evidence across complementary analytical approaches. We identified a reproducible structural-functional network centered on the dmPFC and dACC, which is linked to genes enriched for synaptic function and to serotonergic and dopaminergic receptor profiles. Cortical gradient analysis demonstrated that the aggression network gradient follows the brain’s intrinsic hierarchical organization, with the multimodal gradients significantly predicting its spatial pattern. Overall, this study establishes a unified and reproducible analysis framework that integrates multiscale analysis, thereby providing systematic evidence for the hierarchical brain mechanisms underlying trait aggression.

Based on the comprehensive analysis of whole-brain GMV and functional connectivity network mapping, we revealed a picture of the neural mechanism underlying trait aggression: Trait aggression is significantly correlated with increased GMV in regions such as the dmPFC, ACC, frontal pole, middle temporal gyrus, lateral occipital cortex, cerebellum, and thalamus. This finding is highly convergent with previous research results in the dmPFC and dACC regions (Coccaro et al., 2018; Dugré et al., 2025; Dugré & De Brito, 2024; Seidenbecher et al., 2023). The dmPFC is related to cognitive functions such as metacognition and social cognition, emotion regulation, decision-making, and cognitive control(Kim et al., 2023; Modirrousta & Fellows, 2008); the dACC is related to conflict and error monitoring, effort and cost decision-making, and cognitive control (Shenhav et al., 2016; Vriens et al., 2025). In complex decision-making, the two regions work collaboratively, with dACC monitoring conflicts and dmPFC integrating social value information to form the final choice(Clairis & Lopez-Persem, 2023). This may imply that there are abnormalities in the conflict monitoring, decision-making, and cognitive control functions of aggression. FCNM further showed that these aggression-related brain regions (dmPFC, ACC) are located at the core nodes of the DMN and FPN, forming a pathway from structure to functional network. Recent work suggests that thalamus-centered circuits are involved in bottom-up aggressive behavior, whereas the FPN and DMN are the core networks for top-down control of aggressive behavior intentions(Wang et al., 2024; Wong et al., 2019). This study demonstrates through a series of methods, including self-collected large-scale samples, cross-study, cross-method, and FCNM, that trait aggression encompasses the top-down pathways of the FPN and DMN.

Based on the imaging transcriptomics analysis, we identified a transcriptional component (Component 1) closely related to aggressive traits. This component showed high expression in the ACC and TP regions, and low expression in the occipital cortex. Correspondingly, the upregulated genes were mainly enriched in synaptic signaling-related pathways, involving molecular functions such as monoatomic ion channel activity, cell-cell signaling, and synaptic organization, as well as cellular components such as presynaptic membranes, neuronal projections, and postsynaptic membranes. These genes were significantly associated with inhibitory neurons, excitatory neurons, and astrocytes, and exhibited specific spatiotemporal expression patterns during the critical period of brain development. These results point to emotion-related brain functions. In contrast, the downregulated genes were more heterogeneous, mainly related to visual processing functions. These findings not only confirm the important role of synaptic function integrity in the regulation of aggression (Dugré et al., 2025; Yanowitch & Coccaro, 2011), but also indicate that specific gene expression backgrounds may regulate aggressive behavior by influencing synaptic function and neural network activities(Klasen et al., 2017). From a clinical application perspective, these gene expression patterns may have potential as candidate molecular markers (Xia et al., 2022), and, in particular, ion channel function may represent a potential avenue for future therapeutic investigation.

Receptor distribution analyses further revealed a convergent molecular signature of the trait aggression-related brain network. Most notably, serotonergic (5-HT1A, 5-HT2A, and 5-HT4), dopaminergic (DAT and D1), and noradrenergic (NET) systems showed robust spatial associations across aggression-related brain maps. Although each of these monoaminergic systems has previously been implicated in aggression, they have largely been investigated separately(Fanning & Coccaro, 2015; Zhang et al., 2025). For example, serotonergic signaling, particularly 5-HT2A receptor availability, has been associated with individual differences in aggression in healthy populations (Da Cunha-Bang et al., 2016; Fanning & Coccaro, 2015), while accumulating evidence similarly implicates dopaminergic signaling in aggressive behavior (Dai et al., 2025). Our findings extend these observations by showing that multiple monoaminergic systems converge reproducibly within the same aggression-related neural architecture. This convergence suggests that individual differences in trait aggression may not be associated with a single neurotransmitter system in isolation, but rather with coordinated contributions from multiple neuro-modulatory systems.

From the perspective of the macroscopic organizational pattern of the brain, among the four different modalities of the brain, gradient 1 reflected a transition from single-modality to cross-modality organization, gradient 2 reflected separation along the sensorimotor axis, and gradient 3 reflected separation between the FPN/ventral attention network and the DMN (Huntenburg et al., 2018; Margulies et al., 2016). The spatial pattern of the aggression-related network, as derived from FCNM, was found to align with these findings. Further, the four brain patterns can significantly predict the gradients of the aggression-related networks. Among them, the FCG is the strongest predictor of both gradient 1 and 2, reflecting the core position of the functional organizational pattern in the aggression-related functional network, from perception to cognitive processing. By contrast, the RSG was the strongest predictor of gradient 3, revealing the unique role of the neurotransmitter system in the organization of the aggression-related functional network, and possibly reflecting the neurochemical basis of aggressive behavior.

This study makes three main contributions to understanding the neurobiological architecture of trait aggression. First, we establish a robust structural–functional architecture associated with individual differences in trait aggression. By integrating large-sample individual-level VBM analysis with two complementary coordinate-based meta-analytic approaches (ALE and SDM), we identified convergent structural correlates of trait aggression across individual-level and cross-study evidence. FCNM further extended these structural findings from localized anatomical correlates to distributed functional networks. Together, these convergent analyses move beyond the conventional localization of aggression to isolated brain regions and provide a reproducible network-level framework for further understanding its neurobiological basis. Second, we characterize the molecular signatures associated with aggression-related structural and functional networks. Imaging transcriptomics and neurotransmitter mapping revealed systematic associations with gene-expression patterns related to synaptic function and with serotonergic, dopaminergic, and noradrenergic transporter systems. Importantly, these molecular associations were examined across structural and functional maps derived from complementary analytical pipelines, providing convergent evidence that the macroscale neural architecture associated with trait aggression is embedded within distinct molecular landscapes. Third, we reveal how the aggression-related functional network is embedded within the brain’s intrinsic multimodal hierarchical organization. By integrating gradients derived from morphometric similarity, functional connectivity, transcriptomic similarity, and receptor similarity, we found that the spatial organization of the aggression-related network could be systematically explained by the brain’s intrinsic multimodal gradients. This suggests that the aggression-related network is organized along broader principles of brain architecture spanning structural, functional, transcriptomic, and neurochemical dimensions.

Several limitations should nevertheless be considered. First, the present findings are primarily cross-sectional and correlational and therefore do not establish causal relationships among molecular characteristics, brain network organization, and trait aggression. Second, the molecular findings are based on population-level reference atlases rather than molecular measurements obtained from the participants themselves. In particular, the AHBA is based on postmortem data from a limited number of adult donors, while currently available neurotransmitter maps vary in sample characteristics, spatial coverage, and data quality. Thus, the observed transcriptomic and neurochemical associations should be interpreted as spatial molecular signatures rather than direct evidence of individual-level molecular mechanisms. Third, although extensive sensitivity analyses and complementary analytical pipelines support the robustness of the identified architecture, its generalizability and translational relevance remain to be established in independent populations and clinically relevant samples.

## Data Availability

The multisite structural MRI data, rs-fMRI data, and BWAQ questionnaire data are available on request from the corresponding author; The SDM and ALE data are available at https://osf.io/7nt8m/overview?view_only=50b84fe4058b4f808b9fa60d7c01c5bc; microarray gene expression data are available at the abagen package (https://github.com/rmarkello/abagen); the brain-wide neurotransmitter receptor data are available from the hansen_receptors package(Hansen et al., 2022) (https://github.com/netneurolab/hansen_receptors); the group-averaged functional connectivity matrix from the Human Connectome Project (HCP) is provided in the Brainspace toolbox(Vos de Wael et al., 2020) (https://brainspace.readthedocs.io); we used MRIcroGL(Rorden, 2025) to visualize brain anatomy and BrainNet Viewer(Xia et al., 2013) to visualize glass brains.

## Code Availability

The custom code used in this study is available at https://osf.io/7nt8m/overview?view_only=50b84fe4058b4f808b9fa60d7c01c5bc.

## Declaration of Generative AI Use

We guarantee that AI was not used in any of the core aspects of this manuscript, including conceptualization, data collection, data processing, statistical analysis, and interpretation of results; AI was used solely to assist with language optimization, code checking, literature supplementation, and spelling and grammar checks.

## Ethical statement

This research protocol has been approved by the Ethics Committee on Human Experimentation, Faculty of Psychology, Tianjin Normal University (No. XL202027). All participants provided informed consent and voluntarily participated in the research experiment.

