## supplementary materials for "The multiscale architecture of trait aggression: from meta-analytic convergence to multimodal characterization"

**Appendix 1 Imaging acquisition and preprocessing**

**1.1 Imaging acquisition**

The data for this study were collected from four different sites; the site parameters are listed in Supplementary Table S1.1. The sample sizes for the different sites are shown in Figure S1.1. Since Site 2 contributed the largest sample, we report its collection parameters here: Whole-brain imaging data were acquired using a Siemens 3T Prisma MRI scanner equipped with a 64-channel head coil. Participants lay supine on the scanner bed, and foam padding was placed around the head to minimize head motion. High-resolution T1-weighted structural images were obtained with a magnetization-prepared rapid gradient-echo (MPRAGE) sequence using the following parameters: repetition time (TR) = 2530 ms, echo time (TE) = 2.98 ms, flip angle = 7°, field of view (FOV) = 224 × 256 mm², acquisition matrix = 448 × 512, 192 sagittal slices, slice thickness = 1.0 mm, and voxel size = 0.5 × 0.5 × 1.0 mm³. Parallel imaging (GRAPPA) with an acceleration factor of 2 was employed. Resting-state functional MRI data were collected using a gradient-echo echo-planar imaging (GRE-EPI) sequence with the following parameters: TR = 2000 ms, TE = 30 ms, multiband acceleration factor = 2, flip angle = 90°, FOV = 224 × 224 mm², slice thickness = 2 mm (no gap), and isotropic voxel size = 2 × 2 × 2 mm³. A total of 300 whole-brain volumes were acquired over a period of 10 minutes and 13 seconds. During the resting-state scan, participants were instructed to keep their eyes closed, remain awake, and relax.

**1.2 Anatomical data preprocessing**

Voxel-based morphometry (VBM) analysis was performed using the standard FSL-VBM pipeline (Jenkinson et al., 2012; Smith et al., 2004), consisting of the scripts *fslvbm_1_bet*, *fslvbm_2_template*, and *fslvbm_3_proc*. In brief, T1-weighted (T1w) images were first skull-stripped and segmented into gray matter (GM), white matter (WM), and cerebrospinal fluid (CSF) using the integrated BET and FAST routines within *fslvbm_1_bet*. The resulting GM partial volume images were non-linearly registered to the MNI152 standard space through the creation of a study-specific GM template using *fslvbm_2_template*, which also left-right symmetrized the template. Native GM images were then registered to this template, modulated by the Jacobian determinant of the warp field to preserve local tissue volume, and smoothed with an isotropic Gaussian kernel of sigma = 3 mm (FWHM ≈ 7 mm) using *fslvbm_3_proc*. To mitigate site effects, we applied ComBat harmonization under an empirical Bayes framework (Johnson et al., 2007). These smoothed, modulated GM volume images were then analyzed at the group level using *flameo*.

Cortical surface reconstruction was performed on the same T1w images using the standard recon-all pipeline in FreeSurfer 7.3.2 (RRID:SCR_001847) (Dale et al., 1999; Fischl et al., 1999). The pipeline includes motion correction, intensity normalization (N3 bias field correction), skull stripping, tissue segmentation, and reconstruction of the pial and white matter surfaces. All individual outputs were visually inspected, and manual edits were applied when necessary, following the FreeSurfer guidelines. Cortical thickness was calculated as the distance between the white and pial surfaces at each vertex.

**1.3 Functional data preprocessing**

Resting-state fMRI data were preprocessed using fMRIPrep 23.1.3 (Esteban et al., 2019), a Nipype-based pipeline (Esteban et al., 2019). The main process of this study was based on Wang et al. (2026). The preprocessed data were further denoised using the eXtensible Connectivity Pipeline (XCP) (Ciric et al., 2018). Nuisance regression was performed following the 36-parameter (36P) strategy (Ciric et al., 2017), which includes the six rigid-body motion parameters, mean global signal, mean white matter signal, and mean cerebrospinal fluid signal, together with their temporal derivatives and the quadratic expansions of all terms. Linear trends and intercepts were also included in the regression model. Before regression, AFNI’s 3dDespike was applied to attenuate transient signal outliers, and the first five volumes (dummy scans) were removed to allow for signal equilibration. The regression itself was carried out using linear regression in Nilearn 0.10.0 (Abraham et al., 2014). The residual BOLD time series were then band-pass filtered (0.01–0.1 Hz) and re-censored. Finally, the denoised and filtered BOLD data were spatially smoothed with a 6.0 mm full-width at half-maximum (FWHM) Gaussian kernel. To mitigate site effects, we applied ComBat harmonization under an empirical Bayes framework (Johnson et al., 2007).

**Figure S1.1** Data distribution across different sites

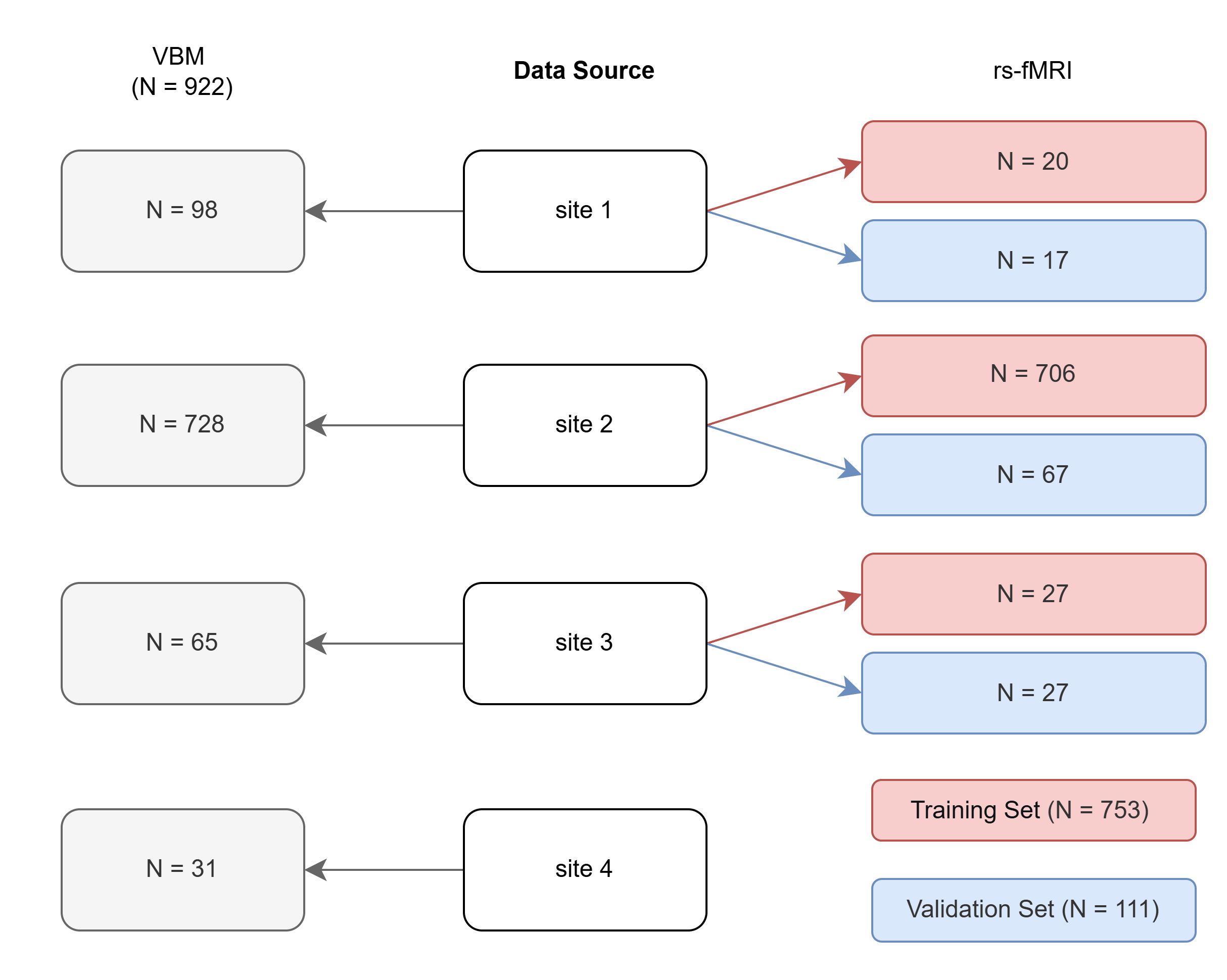

**Table S1.1** Scan parameters for each site

| Site | BNU | TJNU1 | TJNU2 | PKU |
| --- | --- | --- | --- | --- |
|  | 1 | 2 | 3 | 4 |
| Field strength | 3T | 3T | 3T | 3T |
| Slices | 192 | 192 | 240 | 192 |
| Slice thickness (mm) | 1 | 1 | 0.8 | 1 |
| TR/ TE / FA (°) | 2530/2.98/7° | 2530/2.98/7° | 2300/2.07/7° | 2530/2.98/7° |
| Voxel size (mm^3^) | 0.5 × 0.5 × 1.0 | 0.5 × 0.5 × 1.0 | 0.8 × 0.8 × 0.8 | 1.0 × 1.0 × 1.0 |

Notes. FA = flip angle; TJNU, Tianjin Normal University; BNU, Beijing Normal University; PKU, Peking University.

**Appendix 2 Meta-analysis methodology**

**2.1 Eligibility criteria and study selection**

#### 2.1.1 Literature search

A systematic search strategy using two search engines (PubMed and Web of Science) was conducted up to January 14th, 2026, to identify relevant studies. The main focus of the current meta-analysis is on VBM studies, and the following search terms were used: (voxel-based morphometry OR VBM OR structural MRI OR gray matter volume) AND (aggressi*) AND (healthy control* OR healthy volunteer* OR HC). An additional search was conducted by cross-referencing the reference lists of the included articles.

#### 2.1.2 Study selection

Articles were included if they met the following criteria: (1) original study published in a peer-reviewed journal in English; (2) inclusion of a psychometric scale to assess aggression; (3) inclusion of a VBM method for gray matter volume (GMV); (4) conducted a group comparison (or regression) or a dimensional analysis of the severity of aggression; (5) reported peak coordinates and significant effects across the whole-brain (i.e., voxel-wise). Preferred Reporting Items for Systematic Reviews and Meta-Analyses (PRISMA) was followed across the meta-analysis steps.

**2.2 ALE methods and parameters**

We performed a coordinate-based meta-analysis using the GingerALE tool, which implements the activation likelihood estimation (ALE) algorithm (Eickhoff et al., 2009; Turkeltaub et al., 2002). The algorithm models the reported peak coordinates of GMV changes as centers of 3D Gaussian probability distributions, aiming to quantitatively identify topographical convergence across studies while accounting for spatial uncertainty. ALE is a widely adopted method and involves three main steps. First, to compensate for spatial uncertainty arising from inter-template and inter-subject variability in most neuroimaging studies (Eickhoff et al., 2009), all foci are modeled as centers of 3D Gaussian probability distributions. Under the assumption that studies with larger samples provide a more reliable approximation of the “true” location, the distributions are further weighted by the number of participants in each study; thus, larger samples yield narrower Gaussians. Second, the probability distributions of all foci within a single experiment are combined to generate a modeled activation (MA) map (Turkeltaub et al., 2012). Voxel-wise ALE scores are then obtained by computing the union of these MA maps. Finally, to assess statistical significance, the ALE scores are compared against an empirical random spatial null distribution derived from all MA maps. We employed 1,000 permutations and applied threshold-free cluster enhancement (TFCE) for multiple comparisons correction, with a family-wise error corrected significance threshold of *p* < 0.05 (FWE-corrected). Robustness was evaluated using a leave-one-out jackknife analysis.

**2.3 SDM methods and parameters**

We performed a voxel-based meta-analysis using Seed-based *d* Mapping with Permutation of Subject Images (SDM-PSI, v6.23) (Albajes-Eizagirre et al., 2019). This method reconstructs effect-size maps (Hedges’ g) for each study based on the reported peak coordinates and their statistical values (e.g., t-values), allowing inference of meta-analytic effect sizes for GMV changes.

In brief, SDM-PSI first converts the peak coordinates reported in Talairach space to MNI space and extracts the corresponding effect sizes (converting them to t-values if necessary). For peaks with unknown or unreported effect sizes, the software uses MetaNSUE (Albajes-Eizagirre et al., 2019) to estimate the most likely effect size within the bounds of possible effect sizes for each study, and then adds realistic noise, thereby generating multiple imputed effect-size maps. These maps are then combined using a standard random-effects model that accounts for sample size, within-study variability, and between-study heterogeneity (Radua et al., 2012), and the multiple imputations are pooled using Rubin’s rules (Albajes-Eizagirre et al., 2019).

Statistical significance was determined using the threshold typically recommended for SDM meta-analyses, which optimally balances sensitivity and specificity: uncorrected *p* < 0.001 with a cluster extent threshold of 10 voxels (Dugré et al., 2020). Residual heterogeneity across studies was assessed with the *I²* statistic (*I²* > 50% indicating substantial heterogeneity), and potential publication bias was evaluated through meta-regression of effect sizes against their standard errors (i.e., Egger’s test) (Egger et al., 1997; Sterne et al., 2011). In addition, a leave-one-out jackknife analysis was conducted to assess the robustness of the results.

**2.4 SDM sensitivity analysis and publication bias**

Table S2.1 presents the sensitivity analysis and publication bias assessment for the SDM meta-analysis. The results showed low heterogeneity and no evidence of publication bias. Figure S2.1 displays the funnel plot of the significant clusters from the SDM meta-analysis.

**2.5** **Jackknife analysis**

Figure S2.2 presents the overlay of Jackknife T maps for ALE and SDM meta-analysis. The results showed acceptable consistency across ALE and SDM (Table S2.2).

**Figure S2.1** Funnel plots generated by SDM-PSI

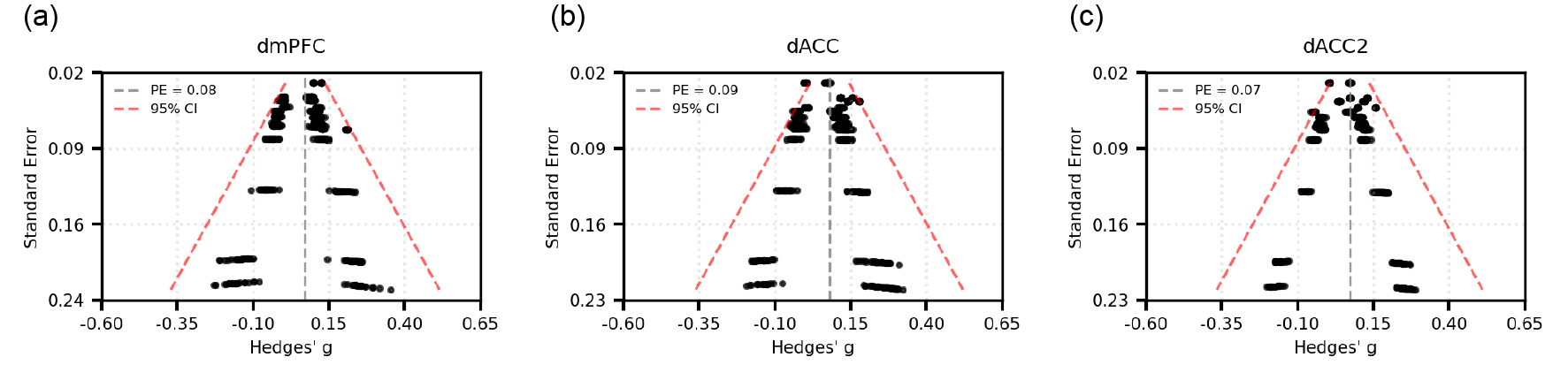

Note. PE, pooled effect (Hedges' *g*); 95% CI, 95% confidence interval.

**Figure S2.2** Jackknife T maps

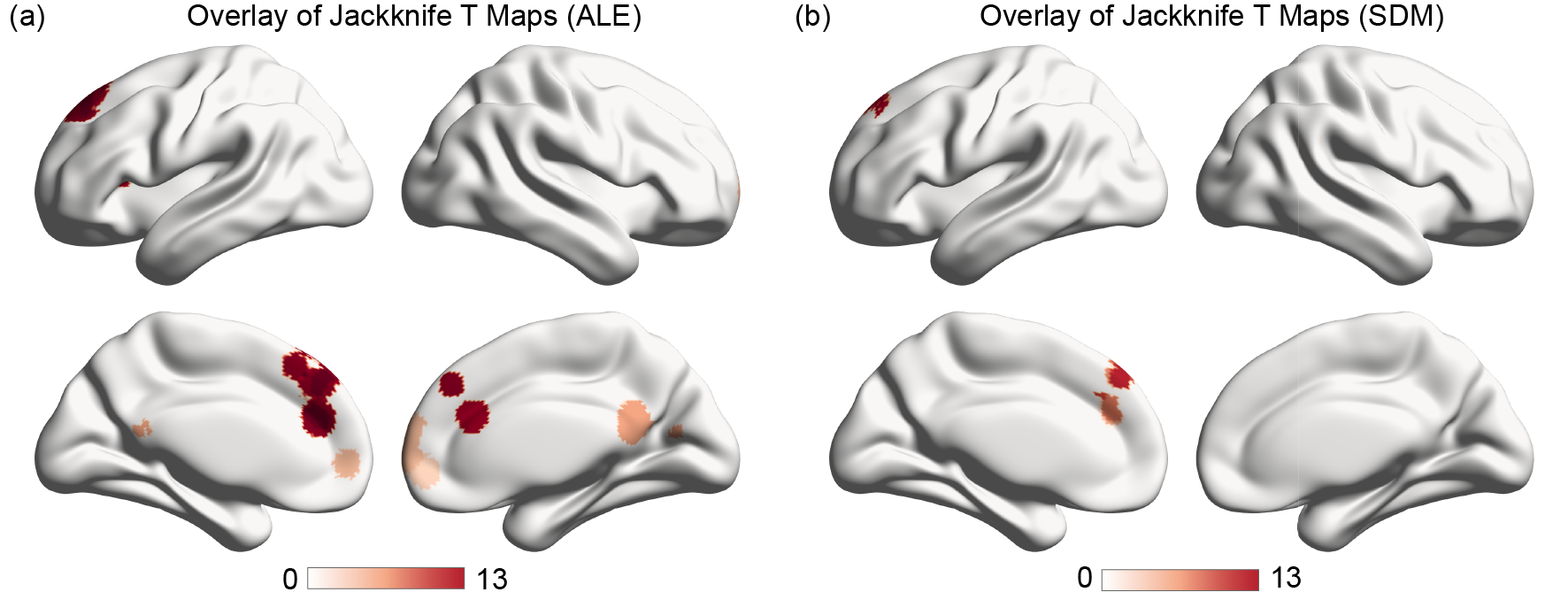

**Table S2.1** The sensitivity analysis and publication bias of the SDM meta-analysis.

| Cluster | Name | Coordinates | *I^2^* | *τ*^2^ | Hedges' g | Metabias test | | | Excess Significance test *p* |
| --- | --- | --- | --- | --- | --- | --- | --- | --- | --- |
|  |  |  |  |  |  | Bias | *Z* | *p* |  |
| 1 | dmPFC | -14,40,42 | 1.88 | <0.001 | 0.08 | -0.37 | -0.42 | 0.674 | 0.668 |
| 2 | dACC | -10,38,28 | 6.88 | <0.001 | 0.09 | -0.17 | -0.19 | 0.850 | 0.387 |
| 3 | dACC2 | -4,32,32 | 1.17 | <0.001 | 0.07 | 0.11 | 0.12 | 0.906 | 0.997 |

**Table S2.2** Jackknife results

| Cluster | Name | Coordinates | SDM | | ALE | |
| --- | --- | --- | --- | --- | --- | --- |
|  |  |  | Overlap quantity | Overlap ratio | Overlap quantity | Overlap ratio |
| 1 | dmPFC | -14,40,42 | 11/13 | 0.85 | 13/13 | 1.00 |
| 2 | dACC | -10,38,28 | 7/13 | 0.54 | 13/13 | 1.00 |
| 3 | dACC | -4,32,32 | 9/13 | 0.69 | - | - |

**Appendix 3 Robustness analysis for functional connectivity network mapping (FCNM)**

To further evaluate the robustness of our main findings, we performed two supplementary analyses. Figure S3.1 illustrates the overlapping patterns of results derived from different seeds, demonstrating consistent spatial convergence across the core regions of the frontoparietal network and the default mode network.

Figure S3.2 shows the significant network obtained from the FCNM analysis applied to an independent validation dataset, along with the overlapping patterns of different seeds. The results of both analyses are consistent with our primary findings, supporting the overall robustness of the study.

**Figure S3.1** The overlapping patterns of different seeds

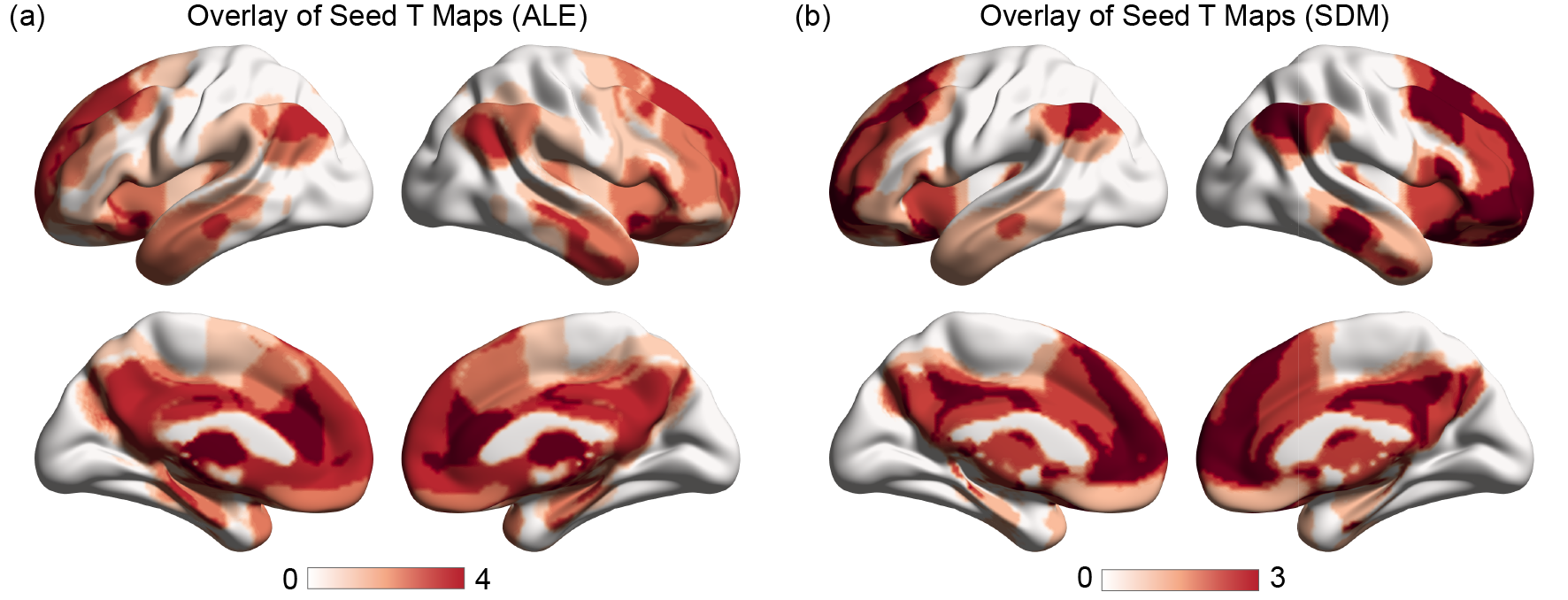

**Figure** **S3.2** The significant FCNM network in the validation dataset and the overlapping patterns across different seeds

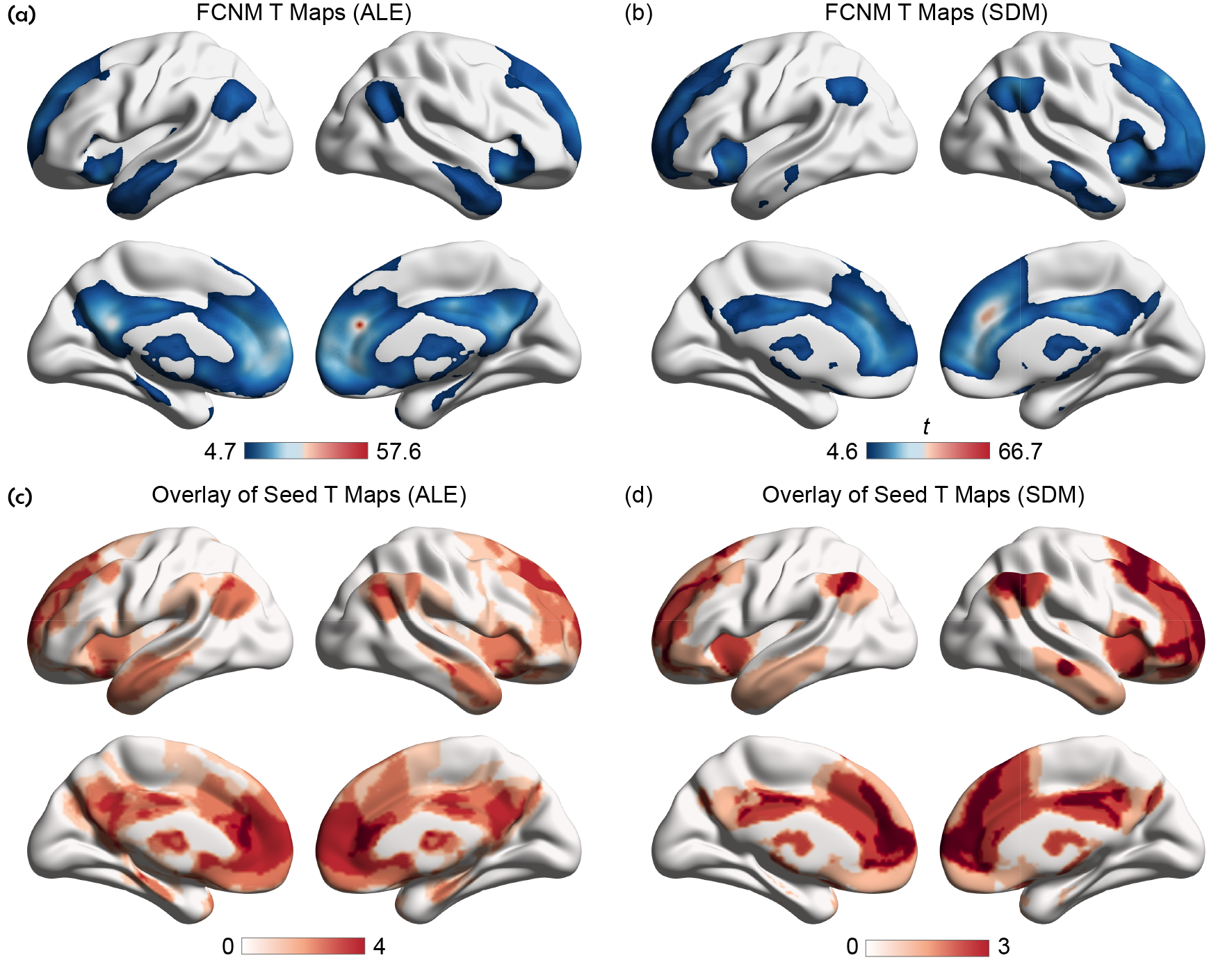

**Appendix 4 Gene analysis**

**Table S4.1** Genes significantly upregulated jointly by ALE & ALE-FCNM

| **Genes significantly upregulated jointly by ALE & ALE-FCNM** | | | | | |
| --- | --- | --- | --- | --- | --- |
| MARCH1 | MARCH8 | ABCB6 | ACADL | ACKR3 | ACOT2 |
| ACP1 | ACTA1 | ACTC1 | ACTG1 | ACTR10 | ADAMTS19 |
| ADAMTS9 | ADAP2 | ADCY2 | ADCY7 | ADGRA3 | ADGRB3 |
| ADGRG1 | ADIRF | ADM5 | ADRA1B | ADRB2 | ADTRP |
| AEBP1 | AHRR | AIFM2 | AKR7L | ALKAL2 | AMIGO2 |
| ANK3 | ANKRD19P | ANKRD50 | ANKRD6 | ANO3 | ANOS1 |
| AP3B2 | AP3S1 | APLN | APLNR | APOM | APOPT1 |
| APTR | AQP4 | ARHGAP33 | ARHGAP4 | ARHGDIG | ARHGEF28 |
| ARMC10 | ARMCX2 | ARPC2 | ARPP19 | ASCL2 | ASS1 |
| ATAT1 | ATOH7 | ATOX1 | ATP2B4 | ATP2C2 | ATP5ME |
| ATP6V1C1 | ATP9A | AZIN1_AS1 | B3GALNT1 | B3GALT6 | B3GAT1 |
| B4GALT2 | B9D1 | BABAM2 | BAIAP2 | BAIAP3 | BATF3 |
| BCKDK | BCL2A1 | BCL7C | BCLAF3 | BCR | BCRP2 |
| BIRC3 | BPHL | BRWD1 | BSPRY | C11orf97 | C12orf45 |
| C16orf58 | C19orf70 | C1QL1 | C1QL3 | C1R | C1S |
| C1orf50 | C20orf27 | C21orf2 | C2CD4C | C2orf69 | C2orf80 |
| C5orf49 | C6orf52 | C8orf46 | CABCOCO1 | CACNA1H | CACNG3 |
| CALHM6 | CALML3 | CAMKV | CANX | CARMIL3 | CASC10 |
| CBLN1 | CCBE1 | CCDC148 | CCDC153 | CCDC167 | CCDC3 |
| CCDC90B | CCKBR | CCNH | CCT5 | CD38 | CD6 |
| CD63 | CD81 | CD83 | CD99 | CDC14A | CDH10 |
| CDH11 | CDH4 | CDH9 | CDKN2D | CDO1 | CEBPA |
| CELF4 | CELF5 | CENPVL3 | CENPW | CEP170B | CERS1 |
| CETN2 | CFAP52 | CFD | CGREF1 | CHCHD6 | CHID1 |
| CHMP1A | CHMP4A | CHST1 | CHST15 | CIB1 | CIB2 |
| CKS1B | CLBA1 | CLPB | CNIH2 | CNN3 | CNR1 |
| CNRIP1 | CNTN1 | CNTNAP3 | COCH | COG1 | COG5 |
| COL21A1 | COMTD1 | CORO1A | CPAMD8 | CPE | CPLX3 |
| CPNE2 | CPNE6 | CPNE7 | CPT1C | CREB3 | CREM |
| CRIP2 | CRMP1 | CTC1 | CTSH | CTXN1 | CXXC4 |
| CXorf57 | CYB5D2 | CYB5R1 | CYHR1 | CYP26A1 | CYP46A1 |
| DACT1 | DACT3 | DAPL1 | DBX2 | DCAF11 | DCUN1D5 |
| DDA1 | DDAH1 | DDAH2 | DDN | DDRGK1 | DHCR7 |
| DHRS2 | DHRS9 | DIAPH3 | DIO2 | DIRAS3 | DKFZp779M0652 |
| DLL3 | DNAAF4 | DNAH14 | DNAH5 | DNAL1 | DNAL4 |
| DNALI1 | DOC2B | DOK1 | DOK6 | DPF1 | DPM2 |
| DPP10 | DPYSL3 | DPYSL4 | DTNBP1 | DYDC2 | DYNLL1 |
| DYRK2 | DYRK3 | E2F3 | E2F5 | ECHDC3 | EDN1 |
| EDNRB | EFCAB1 | EFEMP2 | EFHC2 | EFNB2 | EFNB3 |
| EGFEM1P | EHBP1L1 | ELFN2 | ELL3 | ELOB | EMG1 |
| EMID1 | ENKUR | ENOX1 | EPHA10 | EPOP | ERCC2 |
| ERFE | ERICH6_AS1 | ESD | F12 | FABP5 | FABP5P3 |
| FABP7 | FADS2 | FADS3 | FAM102B | FAM110C | FAM117A |
| FAM120AOS | FAM149A | FAM171B | FAM181A | FAM19A1 | FAM207A |
| FAM229B | FAM71F1 | FAM86B3P | FAM86C1 | FAM86EP | FAM86FP |
| FAR2P1 | FAR2P2 | FARP1 | FBXO2 | FDPS | FGD3 |
| FGGY | FIBCD1 | FILIP1 | FKBP1A | FKBP1B | FOLR2 |
| FOXN3_AS1 | FOXRED2 | FREM3 | FSTL5 | FXYD6 | FZD1 |
| FZD8 | FZR1 | GABBR1 | GABRA3 | GABRA5 | GABRB1 |
| GABRB3 | GALNT17 | GATB | GDA | GGCT | GGN |
| GLDC | GLOD4 | GLRA2 | GLRA3 | GMFB | GMFG |
| GMIP | GNB2 | GNB4 | GNG12 | GNG2 | GNG4 |
| GNPTAB | GOLM1 | GOLT1A | GPC4 | GPM6A | GPM6B |
| GPR143 | GPR68 | GPR88 | GPRIN1 | GPRIN2 | GRIA1 |
| GRID2 | GRIK3 | GRIN3A | GRM3 | GRP | GSKIP |
| GSS | GSTM3 | GTF2H5 | GUK1 | HACD4 | HBQ1 |
| HCG23 | HDGFL3 | HEBP1 | HES1 | HES4 | HES5 |
| HGF | HIST1H2BK | HMGCS1 | HNRNPC | HPCAL1 | HPCAL4 |
| HRAS | HRASLS | HRH1 | HRK | HSD11B1L | HSPB3 |
| HTR1A | HTR7 | HTR7P1 | IDNK | IFI27L2 | IFT22 |
| IGFBP5 | IGSF22 | IL12RB2 | IL13RA2 | IL33 | IMPACT |
| IQCK | ISLR2 | ISYNA1 | JPH3 | JPH4 | JPT1 |
| KCNA5 | KCNG1 | KCNMB2 | KCNMB4 | KCNN3 | KCTD12 |
| KIAA1024 | KIF21B | KIRREL2 | KLF6 | KLF8 | KLHDC8B |
| KLK10 | KLK5 | KLK7 | KRT14 | KRT17 | KXD1 |
| L3MBTL4 | LAMA1 | LARGE1 | LDHA | LDHD | LIN7B |
| LINC00260 | LINC00484 | LINC00900 | LINC00937 | LINC01094 | LINC01140 |
| LINC01197 | LINC01485 | LINC02076 | LINC02217 | LINC02482 | LMAN2L |
| LMO1 | LMO3 | LOC100288911 | LOC100506100 | LOC100507516 | LOC101060091 |
| LOC102724156 | LOC285097 | LOC440934 | LOC441052 | LOC642852 | LOC645166 |
| LOC728392 | LOC728715 | LOC729970 | LOC93622 | LOH12CR2 | LOXL1 |
| LOXL3 | LRRC2 | LRRC36 | LRRC3B | LRRC56 | LRRC73 |
| LRRN1 | LRRN2 | LSM3 | LSM4 | LSM7 | LSP1 |
| LTBP4 | LUZP2 | LY6H | LYN | LYPD1 | LYPD8 |
| LYRM2 | LYRM9 | MACROD2 | MAFG | MAGED1 | MAGEH1 |
| MANEAL | MAOB | MARCKS | MAST3 | MBOAT7 | MCTP1 |
| MCUB | MCUR1 | MDGA1 | ME1 | ME3 | MEA1 |
| MEIS3 | MESP1 | METTL5 | METTL9 | MGAT4B | MGMT |
| MGST1 | MIR7_3HG | MMD | MOB1B | MORN4 | MPV17L2 |
| MR1 | MRAS | MRPL23 | MRPL28 | MRPL37 | MRPS16 |
| MSANTD1 | MSRB2 | MT1B | MT1E | MT1F | MT1G |
| MT1JP | MT1L | MT1M | MT1X | MT2A | MT3 |
| MTA3 | MTCH1 | MUM1L1 | MVD | MYADM | MYDGF |
| MYH7 | MYL12B | MYL5 | MYRIP | MZT2A | MZT2B |
| NADK2 | NAGK | NANOS1 | NAV3 | NBDY | NCAM2 |
| NDUFAF8 | NDUFC2 | NEBL_AS1 | NECAB2 | NEDD4L | NELFE |
| NEURL1B | NFKBIE | NIT2 | NKAIN2 | NKAIN3 | NKAIN4 |
| NKD2 | NLE1 | NMNAT3 | NNAT | NOL4 | NOV |
| NPTXR | NPY1R | NR4A3 | NSFL1C | NT5DC2 | NT5DC3 |
| NTSR1 | NUDT10 | NUDT11 | NUDT14 | NUDT16L1 | NUDT4 |
| NUPR1 | NUS1 | NWD2 | OGG1 | OPRM1 | OST4 |
| OVOL2 | P2RX5 | PACRG | PAK3 | PALM | PAMR1 |
| PAX6 | PCBD1 | PCBP3 | PCDH10 | PCDH17 | PCDH19 |
| PCDH20 | PCED1B | PCSK5 | PDE2A | PDGFC | PDGFRA |
| PDLIM3 | PEA15 | PERM1 | PEX10 | PHPT1 | PID1 |
| PINLYP | PKIA | PLAC9 | PLB1 | PLCB2 | PLPPR2 |
| PLPPR3 | PLPPR4 | PLTP | PMVK | PNCK | PNMA8A |
| PNMT | PNPLA3 | POLE4 | POLR2G | POLR2L | PON2 |
| PPEF1 | PPM1M | PPP4R4 | PPP5D1 | PRKCD | PRKCG |
| PRMT2 | PRMT6 | PRNP | PRR5 | PRRX1 | PRSS23 |
| PRSS35 | PSD3 | PSENEN | PSMB1 | PSMB2 | PSMB7 |
| PSME1 | PSMG3_AS1 | PSORS1C1 | PTER | PTGER3 | PTGER4 |
| PTHLH | PTN | PTPRA | PTPRF | PTRH1 | PTRHD1 |
| PUSL1 | PYCR1 | PYCR3 | PYGL | RAB27B | RAB32 |
| RAB35 | RAB36 | RAB3C | RAET1E_AS1 | RAP2B | RAPGEF4 |
| RAPH1 | RASAL1 | RASGEF1C | RASL10A | RBM8A | RBP4 |
| RCN2 | RER1 | RGMB | RHBDD2 | RIIAD1 | RILPL2 |
| RIMBP2 | RIOX2 | RIPOR2 | RNASEH2C | RNF150 | RNF181 |
| RNF7 | ROBO1 | RPH3AL | RPL12 | RPL31P11 | RPL32 |
| RPL36 | RPL39L | RPL41 | RRAS2 | RRP7A | RRP7BP |
| RSPH9 | RSPO3 | RSPO4 | RTBDN | RTL8A | RTL8C |
| RTN4RL2 | RTP1 | RXRA | RYR1 | RYR3 | S100A13 |
| S100A16 | SARDH | SARNP | SCARA5 | SCG3 | SCGN |
| SCN3B | SCN9A | SCRG1 | SDK2 | SEMA4A | SEMA6B |
| SERTM1 | SFTPD | SGSH | SGSM2 | SH3BGRL3 | SH3PXD2B |
| SH3RF1 | SHANK1 | SHC1 | SHISA5 | SHISA9 | SHISAL1 |
| SKAP2 | SLA | SLC16A2 | SLC26A11 | SLC26A4 | SLC26A4_AS1 |
| SLC27A2 | SLC27A5 | SLC29A4 | SLC30A10 | SLC35F2 | SLC39A12 |
| SLC39A4 | SLC4A3 | SLC7A4 | SLIT1 | SLN | SMAD2 |
| SMAD3 | SMARCD3 | SMIM10L2A | SMIM10L2B | SMIM29 | SMIM4 |
| SNCA | SNHG8 | SNRNP27 | SNX7 | SORBS2 | SOX11 |
| SP5 | SPACA9 | SPATA33 | SPHKAP | SPINT2 | SPON2 |
| SPPL3 | SPRN | SQLE | SSSCA1 | SST | SSTR1 |
| SSTR2 | ST20 | ST5 | STK17A | STK17B | STK25 |
| STK32C | STUM | STX12 | STX17_AS1 | STX1A | SUB1 |
| SULF2 | SUSD1 | SVOP | SYNGR3 | SYNPR | SYT17 |
| TAF10 | TBC1D14 | TBC1D24 | TBC1D26 | TBCB | TCEA3 |
| TDRD9 | TECPR2 | TFPT | TGFBI | THRA | THTPA |
| TIMM10 | TIMM8B | TINCR | TKFC | TLN2 | TM2D3 |
| TMEFF2 | TMEM108 | TMEM120A | TMEM130 | TMEM150C | TMEM158 |
| TMEM159 | TMEM176A | TMEM176B | TMEM208 | TMEM256 | TMEM263 |
| TMEM53 | TMEM54 | TMEM65 | TMSB10 | TNFRSF11A | TNFRSF14 |
| TNIK | TNNI3 | TNNT1 | TOM1L1 | TOX3 | TRAPPC2L |
| TRAPPC6A | TRIM22 | TRIM27 | TRMT112 | TSEN34 | TSPAN33 |
| TSTD1 | TTYH1 | TUBA1B | TUBB | TUBB2A | TUBB6 |
| TUBB8P12 | TUNAR | TXN | TXNL4A | TYW5 | UBA52 |
| UBL5 | UCHL1 | UCHL3 | UFC1 | UG0898H09 | UHMK1 |
| UMPS | UNC5B_AS1 | UQCC2 | VAMP2 | VAT1 | VAT1L |
| VAV1 | VEZT | VIT | VKORC1L1 | VPS37D | VSIG10L |
| VSTM2L | VWCE | WDR54 | WDR6 | WDR66 | WDR86 |
| WFDC1 | WFS1 | WLS | WNT10B | WNT4 | WTIP |
| XKR4 | XYLT1 | YJEFN3 | YPEL1 | YWHAB | YWHAH |
| YWHAZ | ZC3HAV1 | ZCCHC12 | ZCCHC17 | ZCCHC18 | ZFHX3 |
| ZFP36L1 | ZNF226 | ZNF706 | ZNF831 | ZSCAN18 |  |

**Table S4.2** Genes significantly downregulated jointly by ALE & ALE-FCNM

| **Genes significantly downregulated jointly by ALE & ALE-FCNM** | | | | | |
| --- | --- | --- | --- | --- | --- |
| ABCA6 | ABCD2 | ABTB2 | ACAN | ACSL6 | ADAM22 |
| ADAM23 | ADAMTS13 | ADCY9 | ADGRL2 | AGO1 | ALDH1A3 |
| AMMECR1 | ANK1 | ANKH | ANKRD29 | ANKS6 | ARHGEF11 |
| ARID3B | ASB13 | ATG16L1 | ATP13A3 | ATP2B2 | ATP2B3 |
| ATP4A | ATP5MPL | ATRNL1 | BAK1 | BBS7 | BCL3 |
| BCO2 | BEND6 | BICRAL | BMS1P2 | BRINP2 | BUD13 |
| C19orf25 | C1RL | C4orf33 | C5orf24 | C6orf47 | C9orf3 |
| CAB39 | CABP1 | CACNA2D2 | CACNB4 | CACNG8 | CADPS2 |
| CCDC130 | CCDC177 | CCDC186 | CCDC39 | CCDC57 | CCDC58 |
| CCL27 | CCNB1IP1 | CCNO | CDADC1 | CDH12 | CDH7 |
| CDK14 | CDR2L | CELF2 | CENPJ | CEP135 | CEP152 |
| CEP192 | CEP95 | CERK | CFL2 | CKMT1B | CLEC16A |
| CLEC4M | CLGN | CMYA5 | CNST | COG2 | COL27A1 |
| COL5A1 | COL7A1 | CORO6 | CORO7 | COX7A1 | CPLX1 |
| CPLX2 | CREBZF | CSTF2 | CTNNAL1 | CTSC | CUX1 |
| DCAF13P3 | DCBLD2 | DCPS | DDX19B | DEFB131A | DENND1B |
| DENND4C | DENND6A | DEPTOR | DGCR9 | DHX35 | DIS3 |
| DIS3L2 | DMKN | DNAAF2 | DNAJC5G | DND1 | DOK3 |
| DOT1L | DPY19L2P1 | DPY19L2P2 | DSCC1 | DTWD2 | DUS1L |
| ECM1 | EEPD1 | EFR3A | EIF2A | EIF4E1B | EIF5A2 |
| ELMO3 | ELMOD3 | ELOVL4 | EMX1 | ENTPD4 | EPB41 |
| ERCC4 | ERICH5 | ESCO1 | ESRRA | ESRRG | EXOG |
| EXTL2 | EYA4 | FAM110A | FAM120C | FAM135B | FAM13B |
| FAM20A | FAM217B | FAM222B | FAM71E1 | FAM78A | FAM78B |
| FANCI | FASTKD1 | FBN1 | FBXL4 | FBXO32 | FBXO9 |
| FER1L4 | FGF13 | FGF7 | FGF9 | FKBP5 | FLJ23867 |
| FLJ42627 | FNDC5 | FRAT1 | FRAT2 | FRMD4A | FRMD5 |
| FSTL1 | GAS2 | GGA2 | GIT2 | GLCCI1 | GLIS1 |
| GLS2 | GNRH1 | GPAT3 | GPCPD1 | GPLD1 | GPR161 |
| GRAMD1B | GSDMB | GSTT1 | GTF2H2B | GTF3C3 | GUCA2B |
| GUSBP4 | HMCES | HNRNPA1L2 | HR | HS3ST1 | HSD11B1 |
| HSPA4L | HSPBAP1 | HTR1F | ICA1 | IDE | IDS |
| IFNLR1 | IGFALS | IGLL3P | IL17RA | INPP5F | INTS4P1 |
| INTS8 | IP6K2 | IPW | ITFG2 | ITPKC | JDP2 |
| JMJD1C | KBTBD3 | KCNA1 | KCNAB3 | KCNB1 | KCNC1 |
| KCNC3 | KCNC4 | KCNH6 | KCNK12 | KCNT1 | KDM4C |
| KDM7A | KIAA1841 | KIZ | KLF9 | KRIT1 | KRTCAP3 |
| L2HGDH | LAG3 | LCA5 | LEAP2 | LIN28B | LIN9 |
| LINC00515 | LINC00599 | LINC00652 | LINC00999 | LINC01102 | LINC01128 |
| LINC01637 | LINC_PINT | LMLN | LOC100240734 | LOC100286906 | LOC100506124 |
| LOC101928673 | LOC220729 | LOC727896 | LOC729218 | LOC729737 | LPCAT4 |
| LRPAP1 | LRRC4 | LRRC49 | LTB | LURAP1L | LYPD5 |
| LYSMD4 | LYST | MAGI2_AS3 | MAP1LC3B | MAP3K3 | MAP3K6 |
| MAP9 | MASP2 | MBL1P | MBNL2 | MC1R | MCF2 |
| MCM8 | MCTS1 | METTL15 | MFSD9 | MIAT | MID2 |
| MIR124_2HG | MIR29B2CHG | MIR4697HG | MIS18BP1 | MKLN1 | MKNK2 |
| MOGAT1 | MOGAT2 | MORC2_AS1 | MPP1 | MPZL3 | MRPS11 |
| MRRF | MTCL1 | MTPAP | MTUS2 | MYBPHL | MYH7B |
| MYO19 | MYOZ3 | N4BP2L2 | NAA16 | NARS | NCOA3 |
| NCOR2 | NDRG3 | NDUFS1 | NEB | NEXN | NFIC |
| NHLRC2 | NIPAL2 | NKX6_3 | NOM1 | NOXO1 | NR1D2 |
| NR3C1 | NR3C2 | NRG2 | NT5M | NUB1 | OGDHL |
| OSBP2 | OSBPL6 | OSGEPL1 | PAG1 | PANK1 | PATE2 |
| PATJ | PAXIP1 | PCDH7 | PCDHB16 | PCP4L1 | PDE4A |
| PDIA5 | PDZD8 | PECR | PELI3 | PHKA2 | PHYH |
| PI4K2A | PIGC | PIGL | PIM1 | PIM3 | PIN4P1 |
| PINX1 | PJVK | PLA2R1 | PLAGL2 | PLCD4 | PLEKHA8 |
| PLEKHH3 | PLXDC1 | PLXNA4 | POU2F2 | POU3F2 | PPIEL |
| PPIP5K1 | PPP1R16B | PPP1R3E | PRIM1 | PRKCB | PRO1804 |
| PRPF4B | PRRT3 | PSMD6 | PTPN2 | PTX3 | PURA |
| RAD54B | RAPGEF6 | RASSF4 | RELT | RET | RFPL2 |
| RHBDL3 | RICTOR | RMI1 | RMND5A | RNF113A | RNF168 |
| RNLS | RORA | RORB | RPGR | RPS6KA3 | RRM2B |
| RRNAD1 | RSAD1 | RSPO2 | RWDD1 | SCN1A | SCN1B |
| SCRT1 | SDHAF2 | SEC14L5 | SECISBP2 | SEMA4C | SEMA7A |
| SERTAD4 | SERTAD4_AS1 | SGIP1 | SGK494 | SHD | SKIDA1 |
| SLC16A6 | SLC16A7 | SLC17A6 | SLC24A2 | SLC25A37 | SLC38A2 |
| SLC38A9 | SLC4A9 | SLC8A1 | SLC9A5 | SMARCD2 | SNHG1 |
| SNHG14 | SOAT1 | SORBS3 | SPAG4 | SPECC1 | SPNS2 |
| SPTSSB | SRFBP1 | SRSF1 | ST3GAL6 | ST3GAL6_AS1 | STAC2 |
| STARD5 | STARD9 | STIM2 | STOX2 | STPG1 | STRN |
| STS | SUGP2 | SV2C | SYCP2 | SYNE4 | SYT12 |
| SYT15 | SYT2 | TADA2B | TAF1C | TAS2R14 | TBC1D30 |
| TBC1D4 | TBR1 | TDRD1 | TDRD3 | TDRD6 | TENM4 |
| TEPSIN | TFAM | TFB2M | THAP10 | THEM4 | TIAM1 |
| TIFA | TIGD3 | TIMM23B | TMEM127 | TMEM132E | TMEM145 |
| TMEM223 | TMEM81 | TMEM86B | TMTC4 | TPRKB | TPTE2P1 |
| TRANK1 | TRERF1 | TRIP11 | TRPC3 | TRPM3 | TSEN2 |
| TSPAN1 | TTC21B | TTC39B | TTC39C | TUBA8 | TUBD1 |
| UBAP1L | UBE2O | UBE2QL1 | UCP3 | UIMC1 | UPP1 |
| URM1 | USP13 | USP45 | USP47 | VAMP1 | VAV3 |
| VCX | VCX2 | VPS36 | VWA8 | WDR33 | WDR97 |
| WFDC2 | XKR6 | XYLT2 | YBEY | ZADH2 | ZBED5 |
| ZBTB1 | ZBTB38 | ZBTB7A | ZCCHC8 | ZDHHC2 | ZFP14 |
| ZFYVE28 | ZFYVE9 | ZIM2 | ZNF165 | ZNF250 | ZNF333 |
| ZNF385B | ZNF394 | ZNF398 | ZNF491 | ZNF502 | ZNF519 |
| ZNF546 | ZNF670 | ZNF692 | ZNF714 | ZNF718 | ZNF774 |
| ZYX |  |  |  |  |  |

**Table S4.3** Genes significantly upregulated by SDM-FCNM

| **Genes significantly upregulated by SDM-FCNM** | | | | | |
| --- | --- | --- | --- | --- | --- |
| SYT17 | DKFZp779M0652 | ATP2B4 | C1QL3 | TMSB10 | SLIT1 |
| DDA1 | IFT22 | RIPOR2 | EFNB3 | LARGE1 | KIRREL2 |
| EFCAB1 | DYDC2 | CTXN1 | GABRB1 | PKIA | PEA15 |
| MMD | KLK7 | TIMM8B | CHCHD6 | CNIH2 | GABRA5 |
| RIIAD1 | C1QL1 | PCBD1 | POLR2L | RBP4 | STUM |
| PLCB2 | GDA | LY6H | C11orf97 | NUDT11 | TIMM10 |
| WNT10B | PLPPR3 | SMARCD3 | SMIM10L2A | DYNLL1 | PRKCD |
| SNX7 | NTSR1 | LDHD | RAP2B | GABRA3 | CRIP2 |
| DPYSL3 | PRR5 | CHID1 | UBA52 | NECAB2 | SSTR1 |
| PTGER3 | IFI27L2 | SLC26A4_AS1 | C2CD4C | SCN3B | GRIN3A |
| SHISAL1 | PID1 | DDAH1 | PHPT1 | TUBB8P12 | TFPT |
| RRP7A | CEBPA | B3GALT6 | WDR86 | CYB5R1 | PTPRA |
| NKD2 | RNF150 | LOC100288911 | NSFL1C | MYL5 | TXN |
| LRRN2 | CACNA1H | NUDT14 | UQCC2 | RHBDD2 | FXYD6 |
| GSTM3 | ERICH6_AS1 | PTER | BIRC3 | TMEM158 | PPP4R4 |
| SSSCA1 | C5orf49 | DOK6 | LINC01140 | BATF3 | ADRA1B |
| RAB27B | FKBP1B | MGST1 | CREM | CENPW | GNG4 |
| UG0898H09 | KIF21B | TDRD9 | GNB4 | TRAPPC2L | NOV |
| TAF10 | LYPD8 | DNAH14 | ASCL2 | UCHL3 | TMEM159 |
| BAIAP3 | TCEA3 | ARHGDIG | CXorf57 | LINC01197 | ATOH7 |
| KCNMB4 | POLE4 | LRRC3B | CFD | GMFB | TUBB6 |
| NKAIN4 | HPCAL4 | SNCA | TUBB2A | MCUB | FKBP1A |
| TSTD1 | RASAL1 | ZNF831 | ADCY2 | FAM71F1 | YPEL1 |
| PCED1B | SMIM29 | FADS3 | PYCR1 | LINC02217 | GRIA1 |
| CD63 | KLHDC8B | CCDC3 | SH3BGRL3 | CCDC189 | PYCR3 |
| EMG1 | VPS37D | CETN2 | OPRM1 | AP3S1 | CPNE6 |
| HBQ1 | PPM1M | PMVK | PSORS1C1 | APOPT1 | GNG2 |
| PRSS35 | VAV1 | AP2B1 | SLC7A4 | C1S | ANKRD6 |
| SVOP | RAET1E_AS1 | C12orf45 | IL33 | TMEM208 | FAM171B |
| PLPPR2 | FAM120AOS | SLN | MEIS3 | KLK10 | CPNE7 |
| PACRG | SULF2 | WFDC1 | FABP7 | LOC440934 | HRH1 |
| PCDH20 | RILPL2 | GPM6B | NUDT10 | B4GALT2 | MBOAT7 |
| MYADM | DOC2B | MAOB | CENPVL3 | PCSK5 | OST4 |
| ALKAL2 | CNR1 | MRAS | NLE1 | DAPL1 | MRPS33 |
| C8orf46 | FAM149A | RAB36 | RBM8A | PTPRF | UHMK1 |
| CHMP1A | RTN4RL2 | NT5DC3 | KCNA5 | ZNF706 | PCDH17 |
| DDAH2 | WDR66 | RNF181 | SMIM10L2B | ZNF226 | RASGEF1C |
| CDH9 | PRKCG | JPH3 | PNCK | NADK2 | MSANTD1 |
| AMIGO2 | RTBDN | MYH7 | DNAH5 | GGCT | EFHC2 |
| CCBE1 | LSM3 | FAM110C | HIST1H2BK | MZT2A | PPEF1 |
| LMO3 | MESP1 | MANEAL | ZSCAN18 | KCNG1 | B9D1 |
| GMFG | SLA | DTNBP1 | NIT2 | SLC27A5 | EPOP |
| SLC26A4 | PLAC9 | NDUFAF8 | XYLT1 | RIOX2 | SCN9A |
| NELFE | SLC26A11 | CALML3 | WTIP | FAM86C1 | EFNB2 |
| SNRNP27 | TMEM263 | F12 | PSENEN | BPHL | HSD11B1L |
| MT1B | SCARA5 | HTR7P1 | MOB1B | TMEM176A | GLOD4 |
| DYRK3 | COG5 | PDE2A | CACNG3 | COG1 | PDGFC |
| SH3RF1 | TKFC | TBC1D24 | MT1L | MT1G | PNMT |
| MZT2B | ARHGEF28 | ANO3 | MYL12B | NANOS1 | ACADL |
| FDPS | LOC285097 | SST | PEX10 | LRRC36 | SPPL3 |
| BCKDK | PLPPR4 | GUK1 | SFTPD | ICAM5 | TM2D3 |
| MT1E | NKAIN3 | ZCCHC12 | BCR | ENOX1 | HSPB3 |
| LINC02482 | TTYH1 | TMEFF2 | TNNT1 | SLC35F2 | TXNL4A |
| LOC100506100 | CDH10 | ANKRD50 | RER1 | RSPH9 | TUNAR |
| FAM102B | C1orf50 | KCNG3 | SHISA9 | VSIG10L | FZR1 |
| UNC5B_AS1 | MT1X | ETNPPL | SCGN | MAFG | LOC728392 |
| DIAPH3 | NNAT | TOM1L1 | PSMB1 | GPR68 | MACROD2 |
| LOC642852 | CYP26A1 | LSM4 | CHST15 | TMEM130 | LAMA1 |
| FAM207A | HRASLS | E2F3 | EDNRB | TSPAN33 | CNTN1 |
| SPATA33 | BCRP2 | MTCH1 | PTRH1 | DPF1 | CHST1 |
| C19orf70 | E2F5 | FBXO2 | ADTRP | FILIP1 | FAR2P1 |
| PNPLA3 | CDKN2D | TSEN34 | CALHM6 | IGSF22 | SUB1 |
| ADAMTS19 | MARCKS | ANK3 | NBDY | YJEFN3 | DDRGK1 |
| KCTD12 | SNHG8 | MT1JP | MT2A | ME1 | HRK |
| SGSH | CGREF1 | NEBL_AS1 | STX17_AS1 | CD6 | SIL1 |
| HCG23 | GLRA2 | EMID1 | SLC30A10 | ELFN2 | NR4A3 |
| ERFE | HTR1A | ARHGAP28 | EPHA10 | GMIP | SLC4A3 |
| MRPL37 | CAP1 | PSMB7 | RAPGEF4 | CYB5D2 | PAMR1 |
| TMEM54 | CNN3 | TMEM176B | DNAAF4 | SDK2 | MT1A |
| LOC729970 | RPL36 | HCST | NPTXR | CCDC153 | LOC441052 |
| SPINT2 | KLK5 | MPV17L2 | SMAD2 | MR1 | SLC16A2 |
| IL13RA2 | MT3 | ARHGAP33 | MT1F | KCNN3 | DACH2 |
| LDHA | HES4 | GSKIP | HGF | SEMA4F | GAP43 |
| PGM2L1 | AQP4 | S100A13 | LINC00484 | HRAS | RASL10A |
| SMIM4 | PALM | ATP6V1C1 | TMEM256 | VIT | FAM86B3P |
| KCTD4 | CHRDL1 | CIB1 | PYGL | MDGA1 | MTA3 |
| C20orf27 | SYNPR | PUSL1 | LRRC2 | TMEM53 | MCUR1 |
| CMAS | CEP170B | IGF1 | SSTR2 | GTF2F2 | TRIM27 |
| RAPH1 | RNASEH2C | MRPS16 | LINC00260 | GNPTAB | LOXL1 |
| UBL5 | RRP7BP | CHMP4A | PTGER4 | SPACA9 | FREM3 |
| THRA | XKR4 | GPRIN2 | PAX6 | GGN | SLC17A7 |
| ACTC1 | CIB2 | C2orf80 | DZIP3 | SHISA5 | ACOT2 |
| FZD1 | FGGY | CCDC90B | WDR54 | ENKUR | NKAIN2 |
| ASS1 | CDH11 | LOC101060091 | STK17A | MAPK1 | TRMT112 |
| PRSS23 | FSTL5 | RAB3C | ADRA2C | NFKBIE | SYT10 |
| TMEM196 | NOVA2 | RTP1 | YWHAZ | ARHGAP4 | AHRR |
| DYRK2 | FADS2 | CCKBR | CELF5 | SLC39A12 | GATB |
| PTCHD1 | MCTP1 | GLRA3 | VSTM2L | GCNT1 | GDF10 |
| PERM1 | RRAS2 | CD81 | FOXN3_AS1 | BAIAP2 | TUSC3 |
| RPL41 | DNALI1 | TUBB4B | DLG2 | ACKR3 | PRMT2 |
| DCAF11 | ROBO1 | DDN | ARL15 | PRNP | MUM1L1 |
| C6orf52 | RIMBP2 | PNMA8A | PSME1 | MAGEH1 | SUSD1 |
| CANX | KCNK2 | SOX9_AS1 | TLDC1 | CPAMD8 | TNIK |
| MAN1A1 | SCPEP1 | CDH4 | TMEM238 | YWHAH | SOX11 |
| MKL2 | STX12 | VEZT | CPNE5 | TMEM200A | GTF2H5 |
| RPL12 | MAPK3 | LYRM2 | ADGRG1 | RXRA | SLC25A23 |
| CAMKV | RAB35 | NUDT16L1 | DHCR7 | NOL4 | RIPK2 |
| HTR7 | CCDC167 | HACD4 | THTPA | HMGCR | CTC1 |
| ZNF350 | CD83 | GSS | CYP46A1 | RGMB | ADIRF |
| CPE | ST6GALNAC5 | DLL3 | KRT17 | CORO1A | ME3 |
| ANKRD19P | FZD8 | RANBP3L | ACSL4 | CISD3 | SARNP |
| ADAP2 | GRB14 | LYRM9 | TENM3 | VAT1 | NUS1 |
| EXOC6 | SSR4 | GPRIN1 | LUZP2 | MRPL23 | POLR2G |
| MIR99AHG | CERS1 | ARHGAP25 | SLC1A4 | PPP1R32 | PLPBP |
| ELOB | SCG3 | ETFB | SMAD3 | MAGED1 | SH2D5 |
| ZC3HAV1 | PLXNC1 | ECHDC3 | TMEM150C | ACTA1 | ARHGDIA |
| NAV3 | PRAF2 | FAM181A | ACOX3 | KLF8 | ACTG1 |
| ARPP19 | LSM7 | TIMM13 | LOC100129291 | STARD3NL | FABP3 |
| MRPL28 | ZCCHC18 | STK32C | TPST1 | RSPO3 | ARMCX1 |
| MEA1 | OGG1 | ARMC10 | GNG10 | JAKMIP1 | SEMA6B |
| NPY1R | RTL8C | METTL5 | SGSM2 | SUMF1 | LINC01485 |
| CPT1C | MPP6 | ARMCX2 | STX1A | NUDT4 | FAM229B |
| FAM86FP | MIR7_3HG | PDGFRA | SPATC1L | GNG12 | UQCR11 |
| RPH3AL | MRC1 | PCDH19 | IL12RB2 | PTRHD1 | TRIM22 |
| CCT5 | GPR143 | S100A16 | RASL11B | ACTR3B | EFEMP1 |
| ADCY7 | GOLT1A | MYDGF | JPT1 | ASIC2 | FAM213B |
| FTH1 | UBE2Z | WLS | B3GALNT1 | ISLR2 | GRM3 |
| PCDH10 | ATP5ME | NPY | TGFBI | TNFRSF11A | HES5 |
| SEZ6L | MLNR | ADRB2 | FARP1 | DOC2A | DIRAS3 |
| SELENOW | SYN2 | CTSH | GPR150 | IQCA1 | MYRIP |
| MGMT | DNAH2 | EZR | IMPACT | C2orf69 | GBAP1 |
| AZIN1_AS1 | WNT4 | SNAP29 | PSMG3_AS1 | GNB2 | ISYNA1 |
| PYDC1 | SAMD15 | ISOC1 | LOC102724156 | PRRG3 | C1orf194 |
| SKAP2 | GRID2 | SPON2 | NISCH | CHL1 | SLCO1C1 |
| VAT1L | TMTC1 | FDX1L | CMTM8 | DIO2 | PSD3 |
| TSPOAP1 | SPRN | C1QB | ERCC2 | TBC1D26 | WFS1 |
| FGD3 | ARPC2 | TMEM108 | LOC101927027 | NEDD4L | CDC42 |
| COCH | NEUROD6 | FOLR2 | SOWAHA | PLB1 | CNGB1 |
| ARL10 | GABRB3 | FNBP1L | KCTD15 | LRRC56 | JPH4 |
| LRRC73 | FZD9 | SLC17A8 | LINC01094 | HSP90B1 | CAPG |
| ABCB6 | CASC10 | KXD1 | DNAJA4 | NWD2 | LOC645166 |
| UCHL1 | DHX58 | PON2 | KIAA1024 | PSTPIP1 | DENND6B |
| BCL7C | ATP9A | B3GAT1 | ATOX1 | USP9X | FERMT2 |
| TNFRSF14 | HNRNPC | ST20 | ST6GALNAC6 | HGH1 | VWCE |
| GPC4 | TRIM36 | MT1M | PLAC9P1 | LRP1B | GRP |
| ACTR1A | SQLE | ACTR10 | TMEM120A | JUN | TM7SF2 |
| CTNND2 | IL17RD | RGS4 | CCNH | AKR7L | UMPS |
| RAB10 | C1QC | LINC00937 | OVOL2 | CNTNAP3 | CINP |
| TECPR2 | RAVER2 | TMEM47 | COMTD1 | FGF2 | ANKRD30B |
| NPHS1 | FAM212A | NUDT18 | 46083 | GBA | LGI4 |
| BYSL | TMEM107 | CAMK1G | OPRK1 | NT5DC2 | SIN3B |
| 46082 | HMGCS1 | PLTP | STYK1 | GASAL1 | LOH12CR2 |
| MORN4 | CBLN2 | CLPB | MMAA | CDC16 | SYNGR3 |
| CARMIL3 | ANOS1 | PLXNA1 | MYB | SCRG1 | SHANK1 |
| LIN7B | GPM6A | OCA2 | CA10 | WDR17 | CAMK2D |
| UFC1 | PRMT6 | TRIP12 | LYPD1 | NMNAT3 | CLTCL1 |
| ZFP36L1 | ADGRV1 | HPCAL1 | LOC100507516 | CEP83 | PIK3CD |
| FOXG1 | GGTA1P | CDC42BPB | PTPRR | ELL3 | NCAM2 |
| LOR | SNTG2 | ZCCHC17 | KCNMB2 | BTBD9 | FAR2P2 |
| HECTD2 | SLIT3 | MYBPC2 | ENO1 | SLC2A12 | LOC93622 |
| AP3M2 | FAM86EP | MYBPC1 | NEURL1B | SLIRP | LYN |
| FNDC11 | BAIAP2L2 | L3MBTL4 | VSIG4 | NAGK | LOC101927420 |
| ZFP64 | RSU1 | STK25 | C1R | CYBB | CDC14A |
| GRIK2 | C1QA | CHPT1 | MIF | BABAM2 | BRWD1 |
| C19orf12 | TUBA1B | ASL | ALPL | PIANP | MRPS25 |
| LINC00461 | TP53BP1 | DGKB | RPS2P32 | BSPRY | CD38 |
| CMTM4 | LMAN2L | GABBR1 | TSPAN18 | GSPT2 | FCGBP |
| BCAN | DHDH | SERTM1 | GJA1 | TBC1D13 | CYB561 |
| TP53BP2 | DPCD | VKORC1L1 | SPICE1 | FOXRED2 | CORT |
| TPM3P9 | TMEM59L | PPP5D1 | ASMTL | ARHGAP5_AS1 | GPR88 |
| ASXL3 | C16orf58 | RNASEH2A | PHACTR3 | NTNG2 | FZD2 |
| TRAPPC6A | LSP1 | NDUFC2 | WWC1 | PCBP3 | ERC2 |
| GLUD2 | A1BG | TMEM17 | EGFEM1P | EYA2 | CCK |
| PINLYP | HDC | PACS1 | GLUD1 | TBCB | NEK6 |
| WDR31 | CRYM | SPRY1 | GRIK3 | TRIM24 | IFT27 |
| HSPB1 | CCDC68 | P2RY12 | CRIM1 | AK5 | RPL32 |
| TMEM164 | WNT7B | TBXA2R | CSTB | RSPO4 | LINC02076 |
| RAB32 | CARD17 | PGRMC1 | LMO1 | GLT8D2 | ZNF436 |
| MYH15 | FOXF2 | MYL6B | ROMO1 | ALK | NUPR1 |
| APCDD1 | STON2 | ADM5 | SIGIRR | SLC9A6 | HSBP1 |
| DUSP3 | SNTB1 | WDR6 | COL11A1 | SUMO3 | SEMA4A |
| MAPK8IP1 | FAM129B | AK4 | MBOAT2 | MPI | PTGES |
| ZMAT2 | NRCAM | PRRX1 | NEK11 | CD24 | WIF1 |
| PSMB2 | TLN2 | C17orf67 | APOE | BCLAF3 | LRRN1 |
| PCDH8 | KRT31 | KLF10 | NETO2 | PAK3 | TYW5 |
| KRT14 | RTL8A | HDGFL3 | FLRT1 | HES1 | STX7 |
| SH3BGR | GLP2R | MVD | CALU | TSPAN6 | JMJD4 |
| MAST3 | ERI3 | CELF6 | NECTIN3 | ST5 | KIF2A |
| C3orf14 | RYR1 | GPC5 | GABRA2 | PLSCR4 | DNAL1 |
| DACT3 | HDAC9 | ADGRL3 | FAM107A | PON3 | TUBB |
| EFEMP2 | ACOT8 | NUTF2 | RPS20 | B3GNT2 | HEBP1 |
| DHRS2 | CCNYL1 | TTPAL | TINCR | RNF7 | RPL31P11 |
| FAM47E | AIFM2 | ENY2 | DDX10 | C11orf1 | CCDC148 |
| SLC25A22 | DNAJA1 | SIDT2 | GULP1 | ALDH2 | SEC61A2 |
| ASH2L | ACVR2A | TG | CLIC5 | GHR | RPL31 |
| ADGRA3 | TDRKH | NELL2 | TEKT2 | CCR1 | HPCA |
| CXXC4 | NDUFB1 | C16orf95 | CELF4 | LINC01547 | DOK1 |
| DCUN1D5 | PAXBP1_AS1 | SLC2A4RG | HBM | CDC42P3 | PBXIP1 |
| PNPLA4 | YAP1 | ARF3 | PPDPF | UQCC3 | LTBP4 |
| NABP1 | NDUFA11 | LARP1 | FYB1 | COL23A1 | GPNMB |
| CCDC103 | PTRH2 | LYPLAL1 | ADGRB3 | BBOX1 | COL21A1 |
| YBX2 | PTHLH | SGSM1 | LOC728715 | FBXL2 | ADCYAP1R1 |
| CMSS1 | CD248 | LINC00900 | VAMP2 | DHRS9 | SRSF12 |
| SARDH | TNNI3 | S100A6 | IQCK | WISP2 | CTXN3 |
| BORCS7 | CAPZB | CBLN1 | GLI3 | APOM | GPR34 |
| SERP1 | FIGNL2 | THBD | SORBS2 | ADAMTS9 | HSPA13 |
| SRGAP1 | TMEM65 | EDN1 | ASB2 | RCN2 | IFI27L1 |
| KCNN2 | SSC4D | LY86_AS1 | SP5 | CASP1 | SOX5 |
| SIRPA | B4GAT1 | NTSR2 | TCTA | PRDM16 | ZFAND3 |
| CFAP52 | GALNT16 | IL17D | SLC35A2 | DPYSL4 | FAM213A |
| MDFIC | ACAT2 | GALNT14 | CNRIP1 | SORCS2 | CKS1B |
| ASB6 | ALDH9A1 | C2orf73 | CRYZ | OR14I1 | PLPP3 |
| MED16 | TRIB1 | FAM49A | GOLM1 | SELENBP1 | SSBP1 |
| CNIH3 | GGCX | NRP1 | RARA_AS1 | PTN | RUNDC3A |
| SRRD | MFSD12 | PHF13 | MCFD2 | GSTM2 | ID4 |
| CENPF | SLC39A4 | TOMM6 | YTHDF2 | ZRANB3 | PLCH1 |
| LOC101060391 | CC2D2B | DENND2A | DHRS4 | NMT1 | OLFM3 |
| SCAPER | CDO1 | IKBIP | CRMP1 | IGFBP5 | MMAB |
| IDNK | LRRTM4 | GLG1 | SPHKAP | AKAP8 | HEPH |
| P4HA2 | CHST7 | CYP51A1 | SRM | RYR3 | CMIP |
| GPD2 | FIBIN | GAL | RPS24 | ZNF710_AS1 | MDK |
| RGS20 | SMAD1 | NDUFA6_AS1 | TIMP2 | PLIN5 | LOC101927752 |
| RNF11 | MRPL20 | ORC6 | OSBPL8 | NFATC1 | TEX26 |
| PIM2 | TMEM8A | HAUS7 | CDT1 | KLHL25 | UBE2MP1 |
| RLBP1 | C21orf2 | ZDHHC23 | BEGAIN | CASTOR1 | DEAF1 |
| CREB3 | LOXL3 | CDH8 | TYROBP | NAXE | NPDC1 |
| FABP5 | CACUL1 | CLBA1 | NR2E1 | CPLX3 | EIF3J |
| CLDND2 | SKAP1 | ARRB1 | SFTA1P | DBX2 | TAC1 |
| CDC27 | AIF1 | TXLNB | SEMA3D | SH3GLB1 | MAFK |
| SHC2 | TBCA | P2RX5 | ABHD17C | C9orf116 | LONRF2 |
| NDUFB8 | CHCHD5 | S100Z | TMEM117 | SH3GL3 | FAM105A |
| SIDT1 | BHMT2 | ATP2C2 | GSTM5 | RXFP1 | CADM1 |
| ESD | NIPSNAP1 | PGAP1 | CLU | S1PR1 | SWI5 |
| UROS | MSRB2 | ACP1 | C6orf118 | PDLIM3 | FLJ33534 |
| JMJD1C_AS1 | INTU | SHC1 | NIPSNAP3A | UBE2S | LRRC27 |
| FIP1L1 | SLC29A3 | ARHGAP45 | NOL4L | PTS | FAM19A1 |
| TUBB8 | FRA10AC1 | DR1 | CALB1 | HBZ | CLCN6 |
| PITPNM2 | PACSIN1 | MND1 | COL16A1 | NSMCE1 | IDH2 |
| SNCG | AZIN2 | GABARAP | RAB40B | STIM1 | ERBB2 |
| NPPA | TUBB3 | KCNQ1 | LOC105379807 | PRDX4 | THAP8 |
| TMEM141 | RSPH1 | AP1M1 | CASK | PPP1R1A | SLC39A10 |
| GPR6 | LOC100506271 | SRR | SEC24D | LAMC3 | INPP5K |
| DENND1A | SERF2 | RHOJ | CCNA1 | ZC2HC1A | PCDH15 |
| SLC22A18 | RPAP3 | TRMT12 | FAM45BP | POLR2I | FBXL16 |
| LINC00958 | AEBP1 | FSIP1 | TUBA4A | CHRNB1 | PDE1A |
| CYHR1 | YWHAB | PSME2 | PSMB10 | ABHD12B | 46276 |
| ANO2 | SOWAHD | ZMAT1 | LOC105374546 | TEK | MLC1 |
| DMAC1 | CABYR | STK17B | KLHL35 | 46089 | SHISA4 |
| RPS18 | MAPK11 | COL5A3 | CBFA2T3 | CTSO | ARL16 |
| ZBTB44 | P2RY13 | TMEM234 | RARRES3 | DENND1C | THNSL2 |
| VPS29 | RETREG1 | SAR1A | AKR7A2 | CNPY2 | OTUB2 |
| DISP2 | GJB6 | PHF24 | LINC00319 | CCDC71L | PUF60 |
| TAMM41 | DKK3 | EIF4EBP1 | DPP10 | BTC | FXYD1 |
| HEYL | NEXMIF | DUSP6 | LAMTOR4 | NCAM1 | ARHGAP36 |
| CHST9 | UBE2B | ROBO2 | MGAT4B | BCL11B | BCL2A1 |
| CCL28 | GALM | GNAQ | PHF20L1 | ARFGEF3 | CTH |
| GPR22 | RAB31 | HCFC1R1 | CHMP2A | KRT83 | RILPL1 |
| VBP1 | TRIB3 | DAD1 | AGA | VPS26B | CHD3 |
| PKIB | RSPH14 | ADAMTSL2 | HYI | ZNF727 | DDIT4L |
| DIAPH2 | FAM241B | GTDC1 | DUSP18 | PPIL6 | TBC1D14 |
| ADORA2B | CCDC85A | PTGR2 | FAM174B | PTDSS1 | UBE2D2 |
| FABP5P3 | LCLAT1 | DCTPP1 | FAM117A | SOBP | ANGPTL4 |
| PRPH2 | CARD16 | ZFHX3 | SLC29A4 | DPP10_AS1 | LOC103344931 |
| PRICKLE2 | C1orf52 | EHBP1L1 | FIBCD1 | GYG2 | TRABD2A |
| CDH13 | GIT1 | EMC7 | ACSS3 | NCAN | ARHGAP42 |
| CYTOR | LINC01551 | MANF | SPATA18 | CACHD1 | KCNB2 |
| CCDC85C | PCDHB4 | MSMO1 | KCNT2 | HERC5 | OAF |
| GLDC | TYRO3 | GALNT17 | PPP1R9B | SLC27A3 | CNMD |
| FFAR4 | DUSP2 | ZNF229 | CLDN10 | DNAJC2 | SERPINI1 |
| ATL1 | ACTG1P4 | MRPL41 | RMI2 | KLHDC8A | KPNA2 |
| PRELP | C6orf62 | LOC105372480 | TMSB4X | GXYLT2 | STOX1 |
| A1BG_AS1 | DOK4 | C16orf74 | C12orf73 | METTL6 | EIF1 |
| ANKLE2 | CAMLG | RPL27A | DMGDH | SRI | MRPS21 |
| ARPC3 | CNTN6 | APBA1 | PXMP4 | SLC8A2 | SLC44A2 |
| PACRGL | ABHD1 | SMKR1 | MAPK8IP1P2 | JADE2 | DNAL4 |
| GRK6 | SOX9 | COMT | TSC1 | GRM8 | ACOX2 |
| APEX2 | C10orf82 | FUOM | GSTM1 | FAS | WIZ |
| VPS37B | COPZ1 | CDC42SE2 | TAF9B | ME2 | SERHL2 |
| RPL38 | PRR29 | NDP | CHODL | WDR25 | COPRS |
| FAH | NUPR2 | MPPED1 | ANO10 | MAPKAP1 | GOLGB1 |
| PDZD11 | SELENOM | P2RY1 | RAN | EPHX1 | KLHL12 |
| LKAAEAR1 | NSMF | COLEC12 | SCN3A | BAALC | COPS4 |
| HCN4 | POC1A | SNX10 | CDYL2 | ATAT1 | C12orf29 |
| MEIG1 | TOLLIP | HHIPL1 | FGFBP3 | TTPA | SCN7A |
| ISG15 | SIVA1 | PEMT | HTR2C | RNF43 | FAT1 |
| SLC16A9 | NWD1 | BAG2 | NUMBL | FHOD3 | DRAM1 |
| LRRC42 | TMEM258 | FAU | EMP3 | SSR4P1 | CCDC158 |
| TCERG1L | KCTD21 | C16orf89 | STBD1 | TRUB1 | PAPLN |
| PDGFB | FADS1 | PPP2R5C | LINC02361 | DHCR24 | SYT5 |
| C16orf87 | TMEM42 | CABCOCO1 | NHP2 | METTL9 | 46082 |
| CRLF1 | PTP4A3 | RBBP7 | RAP1GAP | FAM171A2 | RGS22 |
| PGAM5 | ATXN10 | RASA4CP | ADGRF5 | SC5D | DACT1 |
| LOC102723665 | BAD | GPX4 | CAPN12 | DOCK7 | CFAP44 |
| TNFAIP8L3 | DENR | NR1H3 | BEX3 | SAMD14 | NPY5R |
| SRP19 | EIF6 | MEST | CDHR1 | CCNY | ASTN1 |
| GPR26 | GRASP | LARS2 | DRAIC | ARL8A | ALDH6A1 |
| LIX1 | CHDH | TSPAN4 | LINC01750 | PUM1 | CACNB2 |
| FBXW9 | F3 | KLF6 | GDPD2 | CERS6 | PPP2R5A |
| RITA1 | NPM3 | SLC40A1 | RPL27 | GUF1 | HMGB3 |
| AP3B2 | YKT6 | APLNR | CSGALNACT1 | SNX3 | SYNM |
| NACC1 | WNT10A | RNF182 | KLRC2 | NXPH1 | EDARADD |
| SUSD4 | TOX3 | RPL29 | CELF3 | PNOC | TNFAIP8L1 |
| FUCA1 | MARCO | C19orf53 | ARHGDIB | CPNE8 | MORN3 |
| RASGRP1 | ZIC5 | VPS35 | RNF157_AS1 | TADA3 | BEX2 |
| SCUBE2 | MRPL55 | HBG1 | GAMT | DNAJC24 | SERPINI2 |
| TRPS1 | RRAGB | 46268 | TLR3 | CCDC149 | C12orf75 |
| SNAPC5 | TPI1 | PPM1N | GNAZ | TPD52 | VANGL2 |
| SLC1A2 | ADCYAP1 | MAGIX | SBF1 | ACTB | TMEM18 |
| CA11 | SCG2 | LIMS2 | APTR | CNTN3 | ELMO2 |
| SMIM24 | COMMD9 | FARSB | FTL | ERICH1 | RPL39L |
| PET100 | NDUFA3 | ALKBH7 | CCDC112 | WDFY4 | MIF_AS1 |
| SPRY4 | ICAM3 | GPR27 | CCDC8 | AHNAK | B4GALT3 |
| LHX2 | SLC25A18 | PPP1R3G | SERINC3 | GABRG1 | COLCA1 |
| LOC101930085 | S1PR2 | C2CD2L | LOC100130331 | TSPAN2 | ALCAM |
| MED8 | GRIN2B | C1orf122 | TMEM136 | ITGB8 | NAA38 |
| JAG2 | GPT2 | ENHO | STMN1 | ZFP69 | C12orf76 |
| SLC19A1 | CCDC28A | RDH11 | NECTIN2 | VASN | CENPP |
| SCML1 | NSG2 | REEP5 | GLI1 | TMEM233 | PPP1R21 |
| MRPL44 | NXN | PKP3 | ADAMTS7 | BDH2 | TRAPPC5 |
| TMPRSS5 | ZZZ3 | SPRED1 | PRKG1 | WDR13 | B3GNTL1 |
| ZFYVE1 | MPHOSPH6 | DNAH7 | LIX1L | SIX3 | USP35 |
| RPL24 | WDR7 | HINT2 | SEC14L2 | ARHGEF40 | MARK1 |
| PTCH1 | RAC2 | RGS10 | ALG1L | SH3PXD2B | TMEM200C |
| MARK4 | DCN | DPM2 | NMU | NCKAP1 | NDST3 |
| TIMP3 | ERLIN1 | RDH10 | LNPK | DHRSX | PIRT |
| SYTL5 | SLC15A2 | TUBA4B | DERL2 | DCLK2 | SYTL2 |
| TEX30 | RTN4RL1 | CCND3 | TBPL1 | DYNC1I1 | SEM1 |
| HMGCL | FERMT1 | PMM1 | POMGNT2 | SOX3 | C1orf115 |
| NYNRIN | NUDT2 | BEX1 | PLXNA3 | SMIM30 | OAZ2 |
| BTNL9 | ARPC5 | HSPB11 | SYTL4 | FAM189A2 | LSAMP |
| ENTPD6 | PROCA1 | MAP1LC3A | SHISA7 | LOC105374952 | CLPS |
| SIRT3 | PARD3B | SNAI2 | NAA10 | APBB1IP | RALGPS2 |
| CHPF2 | PPP1R26 | KIF3C | ZNF277 | LFNG | UBE2E2 |
| DUSP28 | SLC8A3 | WSB2 | CPNE2 | KIAA1328 | SLC27A2 |
| B3GALT2 | DLK2 | EMC3 | GPC1 | PDGFA | KIAA0319 |
| FAM131A | TMEM132A | SNX2 | CDC42EP4 | ATP5MF | RCN1 |
| EFHB | RGS18 | OPHN1 | CACNG4 | OLFML2B | AGAP3 |
| AKR1C3 | PIFO | DGCR6 | GADD45GIP1 | CRYZL1 | ZMYND10 |
| NENF | PSMA6 | RREB1 | HIST1H2BC | NGB | BLVRB |
| LRRC61 | CLINT1 | ZNF521 | SPATA20 | SLC2A10 | TMOD3 |
| TSKU | CLEC9A | TIMM50 | ABCC6 | SLC7A11 | ANKRD30BP2 |
| CSPG5 | DMD | HSP90AB1 | P2RY6 | PCDHGC3 | KRT86 |
| NEUROD2 | ZMYM3 | CXCR4 | CHRM3 | CHST8 | CX3CR1 |
| RBP1 | LGALS1 | ABR | FAM83H | HHLA3 | SERPINB2 |
| SNTA1 | CXADR | ARNT2 | SOX4 | NETO1 | TP53TG1 |
| TPT1 | MEDAG | CD99 | DCAF15 | SMIM27 | FUT9 |
| RIMS1 | MAFF | GSTO2 | ITGB4 | HSDL2 | HSDL1 |
| CASQ1 | NUP85 | RFC3 | ONECUT1 | LOC100507351 | OAZ3 |
| MINOS1 | RAB34 | GPX3 | SLC44A3 | LAMTOR2 | RARS |
| AKAP5 | SAP30BP | RPL8 | LPP | GFER | MRPS6 |
| APLN | USE1 | GRAMD4 | TPPP3 | ZNF572 | NTAN1 |
| MCM6 | CYS1 | SYN1 | NDUFAF4P1 | PHGR1 | TNR |
| ZNF646 | FCGRT | PARP1 | APRT | TSTA3 | TIMP1 |

**Table S4.4** Genes significantly downregulated by SDM-FCNM

| **Genes significantly downregulated by SDM-FCNM** | | | | | |
| --- | --- | --- | --- | --- | --- |
| NCOA3 | TRPC3 | MIR29B2CHG | URM1 | SYCP2 | MAGI2_AS3 |
| LYSMD4 | ANK1 | SLC16A7 | RPGR | IFNLR1 | SYNE4 |
| SNHG14 | HS3ST1 | GPCPD1 | CUX1 | EIF2A | LOC100506124 |
| CENPJ | FASTKD1 | GPLD1 | KCNAB3 | EEPD1 | GLCCI1 |
| ST3GAL6 | DIS3L2 | STARD5 | INTS4P1 | RORA | LINC01637 |
| CKMT1B | UPP1 | LRRC49 | TMEM86B | AGO1 | MC1R |
| EPB41 | SLC25A37 | SEMA7A | PLA2R1 | KCNK12 | SCRT1 |
| PRO1804 | IDE | TTC21B | UCP3 | KCNC3 | CLGN |
| SECISBP2 | L2HGDH | TAS2R14 | CCDC58 | ESRRG | SUGP2 |
| ENTPD4 | ZYX | TMEM145 | SLC38A2 | NT5M | CEP95 |
| ZBTB1 | FRAT1 | USP13 | MORC2_AS1 | CDH7 | IGLL3P |
| FGF9 | NRG2 | C19orf25 | IDS | SGK494 | LAG3 |
| ZADH2 | SCN1B | CACNA2D2 | RHBDL3 | CEP152 | THAP10 |
| C4orf33 | ASB13 | GPAT3 | OSBP2 | CLEC16A | ADAMTS13 |
| LRCH1 | CDR2L | RSPO2 | TUBD1 | TDRD6 | LOC220729 |
| KCNC1 | ATP2B3 | ST3GAL6_AS1 | JDP2 | NEXN | NFIC |
| TFAM | PPIEL | SERTAD4_AS1 | FNDC5 | TIFA | LRPAP1 |
| CDADC1 | STRN | PIGL | KLF9 | COL5A1 | DCBLD2 |
| SGIP1 | BAK1 | TRANK1 | MKNK2 | PAG1 | NR2C1 |
| RAD54B | KREMEN1 | OSBPL6 | GNRH1 | RPS6KA3 | CAPN15 |
| UIMC1 | GUCA2B | ZNF333 | COL7A1 | ANKH | DTWD2 |
| SLC4A9 | SYT2 | FBXL4 | KCNA1 | LINC00599 | SRSF1 |
| CREBZF | METTL15 | DCPS | AMMECR1 | FAM71E1 | ZNF268 |
| FBXO9 | CADPS2 | RMND5A | ACSL6 | DEFB131A | ANKS6 |
| PLCD4 | TMEM132E | RET | CACNB4 | ATP8B2 | CCNB1IP1 |
| C9orf3 | FSTL1 | GTF3C3 | ZCCHC8 | VAV3 | UBE2QL1 |
| LIN9 | ZFYVE28 | PECR | SNHG1 | ELMOD3 | ACAN |
| ZNF718 | PIM1 | KBTBD3 | ITPKC | STARD9 | IGLL1 |
| KRIT1 | SORBS3 | LINC00476 | INTS8 | UBAP1L | DPY19L2P1 |
| LPCAT4 | FAM110A | PIF1 | TLE2 | ITFG2 | SPTB |
| TAF1C | ZNF502 | FBN1 | LCA5 | LOC100240734 | METTL24 |
| USP45 | CMYA5 | GAS2 | KRTCAP3 | BUD13 | MIAT |
| 46277 | SLC16A6 | CCDC177 | ZBED5 | PATJ | SCN1A |
| RAB37 | LYPD5 | XKR6 | RMI1 | PLK5 | WDR90 |
| IPW | FBXO32 | LRRC37A4P | NEIL1 | MGARP | PLXNA4 |
| PTPN2 | ZNF37BP | RWDD1 | RNF113A | GIT2 | NKX6_3 |
| SERTAD4 | STPG1 | DCUN1D2 | GTF2H2B | TBC1D4 | DENND1B |
| OSGEPL1 | ATP13A3 | DIS3 | FRAT2 | MTPAP | EXTL2 |
| SPAG4 | ZSCAN9 | LINC01102 | GPATCH8 | FER1L4 | ADAM22 |
| TTC39B | KCNC4 | FAM214A | C6orf47 | MTCL1 | ZNF670 |
| GSTT1 | DSCC1 | GPR161 | VCX2 | LURAP1L | FLJ42627 |
| RRNAD1 | DND1 | AHI1 | CACNA1A | HSPA4L | COG2 |
| NFKB1 | TARDBP | ZZEF1 | ZNF692 | ITM2A | CPLX1 |
| GNB3 | JAK2 | ARHGAP9 | LOC401357 | VCX | EXOG |
| MCTS1 | PVALB | GUSBP4 | TSEN2 | TRERF1 | ZNF362 |
| BICRAL | SCLT1 | ZNF546 | TBCE | DHX35 | GSDMB |
| ZIM2 | CLMP | MYH7B | CSTF2 | MIS18BP1 | NR3C1 |
| CARD9 | STARD10 | CYB5R4 | VAMP1 | TPTE2P1 | GGA2 |
| NAA16 | FAM131B | HTR1F | ESRRA | PIGC | DACT2 |
| ABCA6 | N4BP2L2 | DOT1L | GIGYF2 | SHD | RNF165 |
| PLXDC1 | RNLS | PRIM1 | POU2F2 | HMCES | NFATC2IP |
| SPTSSB | ADCY9 | SMCO4 | RASSF4 | SLC38A11 | EIF5A2 |
| ERP44 | OVGP1 | IL17RA | TWNK | SLC17A6 | PLEKHA8 |
| RSAD1 | SNRNP35 | BEND6 | B3GALNT2 | MAP3K6 | PKD1P6_NPIPP1 |
| FAM13B | LINC01137 | MYO19 | LOC101928673 | DDX11 | CNOT6L |
| FLT3 | TTC39C | PDIA5 | ATP4A | TRIP11 | CRABP1 |
| GLS2 | ARMCX4 | SKIDA1 | ZNF557 | HERC3 | TCTEX1D1 |
| MIR124_2HG | FAM43A | ZFP14 | NEB | POMC | BRINP1 |
| MOGAT1 | FOXN3 | SMARCD2 | PAXIP1 | GLIS1 | CCDC57 |
| ERICH5 | ANKRD29 | DMKN | JMJD1C | MLXIP | ERCC4 |
| DPY19L1P1 | PTX3 | SEMA4C | TMEM127 | LOC729737 | CEP192 |
| CLK3 | MPZL3 | ABCC5 | RBBP4 | PLEKHH3 | PPP1R16B |
| FAM20A | TEPSIN | FRMD4A | SLC9A5 | LOC100294145 | TRIM52 |
| PLCXD1 | MCF2L | MTRF1 | SLC5A6 | RAB24 | SIRT4 |
| RSRC1 | PPFIA4 | PHYH | GART | ICA1 | STAC2 |
| STS | MBNL2 | HR | PCDH7 | CHST12 | LINC00515 |
| ADD1 | NR3C2 | EMSY | CEP85L | LEAP2 | CCL27 |
| FAM222B | POU6F1 | PPARD | MOGAT2 | ECM1 | FGF13 |
| ADM | PRPF4B | DPY19L2P2 | LMLN | PATE2 | TPRKB |
| ZBTB38 | NDRG3 | BRWD3 | LINC00265 | MSRB3 | NOXO1 |
| CCDC88B | ADAM23 | DIDO1 | IGFALS | LRRC4 | NUB1 |
| POU3F2 | DNAJC25 | ZNF774 | STIM2 | CDS1 | TGM1 |
| CTNNAL1 | PJVK | C15orf59 | TMTC4 | MRGPRF | RORB |
| WDR97 | ATRNL1 | DUS1L | SERPINB9 | JADE1 | ARHGEF18 |
| CTSC | DENND4C | LINC_PINT | CORO7 | TNIP1 | SRFBP1 |
| ATG16L1 | SRSF11 | USP28 | SRPK1 | NCOR2 | SLC24A2 |
| NME5 | FLJ23867 | EYA4 | FAM216A | PPIL3 | IP6K2 |
| SOAT1 | HAUS8 | PTPRD_AS1 | DENND6A | KDM7A | RANBP17 |
| CEACAM19 | ESCO1 | CABP1 | SEC31B | SLC35F3 | MPP1 |
| LOC100130950 | CCNO | SLITRK3 | RAPGEF6 | BBS7 | CROCCP2 |
| WDR33 | FAM133DP | INPP5F | PCNT | CEP135 | NHLRC2 |
| PKD2 | LOC729218 | VWA8 | NAALAD2 | MRS2 | KBTBD2 |
| TRIM7 | P3H3 | SMIM3 | C1RL | PANK1 | ANKS3 |
| MASP2 | AR | TCERG1 | YTHDC1 | RICTOR | PIM3 |
| RGS6 | ABCD2 | SLC39A13 | ZDHHC2 | TBC1D16 | PDE4A |
| ZNF250 | UPF3A | VPS54 | DDX46 | MCF2L_AS1 | PHKA2 |
| SLC35A4 | TRMT2B | SRSF8 | CTIF | C4orf3 | TSPAN9 |
| MAP3K3 | LAT | TDRD3 | RELT | GRAMD1B | LOC389834 |
| HSBP1L1 | ATP5MPL | TIAM1 | ARID3B | PPIP5K1 | SV2C |
| PIN4P1 | NR1D2 | TRMT9B | EFR3A | CCDC130 | KRTAP5_2 |
| MAP1LC3B | LOC101928087 | TNF | VIPR2 | LOC727896 | TFB2M |
| COL13A1 | MCF2 | RDH5 | TOB1_AS1 | ALDH1A3 | NPIPA8 |
| ARHGEF11 | ADAMTS5 | STRBP | TDRD1 | FANCI | ATP2B2 |
| KCTD9 | LAMA3 | PDP2 | SYT12 | KCNA2 | SRRM3 |
| STOX2 | LYST | PPARGC1A | QRFPR | XYLT2 | ADAM8 |
| TNFRSF25 | TENM4 | KLF12 | KDM4C | DNAAF2 | ELOVL4 |
| RBM14 | KLHL20 | PDZD8 | TMEM81 | ZFYVE19 | TAS2R10 |
| CDK13 | ZNF398 | SLC9A1 | MTUS2 | MIR4697HG | TADA2B |
| SETD7 | TRIM26 | ZNF273 | DCP1A | CERK | PCP4L1 |
| RBAK | PPP1R15B | MRRF | MGST2 | CCR10 | BMS1P1 |
| MED11 | PRR16 | SPTBN4 | SSH3 | SRA1 | HIST1H4A |
| AAAS | LRR1 | CHRD | NIPAL2 | CDH12 | RASD2 |
| SCN4B | BTBD3 | KALRN | CCDC186 | ZNF519 | ARID2 |
| KBTBD12 | ZNF385A | PKD1 | VPS36 | ZNF165 | OLR1 |
| DNAJC4 | HPS1 | TBC1D30 | COL27A1 | BMS1P2 | AP5B1 |
| MIR600HG | LINC00963 | SYN3 | PRMT7 | COX7A1 | DPY19L2P3 |
| TRMT61B | RELL2 | FAM49B | MFSD9 | LEO1 | BHLHE40 |
| SOCS5 | LOC652276 | EPN3 | HIST1H1D | MAP9 | LIN52 |
| IFRD2 | FADS6 | USP36 | ALX3 | SPART | PLEKHM2 |
| PHC2 | PATL1 | ANKS1A | CASP8AP2 | NAAA | ZNF394 |
| SPNS2 | SEC14L1 | PKD2L2 | PINX1 | TAS2R13 | NOP53 |
| SETD6 | BHLHE41 | HYLS1 | PLAGL2 | CPSF1 | TIMM23B |
| SLC38A9 | NOM1 | TUBE1 | STAG3L4 | RALGAPA2 | MAN2A2 |
| CDKN1B | TIA1 | FAM120C | C5orf24 | INSM2 | PDP1 |
| PRSS56 | RUNDC3B | ARMT1 | PASK | DDX19B | SMC2 |
| TCHP | VWC2 | ANKRD16 | EIF4E1B | DVL2 | BCO2 |
| ZMYND8 | EEF2K | PCGF1 | IDI2_AS1 | NSMAF | ZMAT4 |
| PELI3 | ABTB2 | KDF1 | MICU3 | BSDC1 | KCNH6 |
| MTDH | ZNF630 | USP47 | ADCY3 | ZNF786 | ZNF383 |
| TPTE2P6 | PLCB4 | DCAF13P3 | KIAA1841 | DNAJC5G | KIAA1614 |
| PENK | HNRNPA1L2 | SLC39A8 | CELF2 | MYBPHL | DENND4B |
| CCDC136 | LXN | LINC00999 | ANKRD26 | PNISR | EYA3 |
| FAM76B | TMCO6 | WFDC2 | PHAX | TIGD3 | WDR19 |
| NUP54 | SLC2A11 | THEM4 | TMEM237 | PPP1R3E | PSMD6 |
| MYOZ3 | CCDC152 | SLC19A2 | CROCCP3 | STRIP2 | PDXDC2P_NPIPB14P |
| TBR1 | SLC25A12 | MCM3AP_AS1 | CAMTA1 | PURA | PLAGL1 |
| OGFOD2 | NUDT5 | KRBA1 | CFAP65 | USP49 | ITGB3BP |
| TAS2R30 | STH | RBM41 | FAM217B | FBXL12 | C2CD3 |
| ATF6B | KLF13 | CCDC146 | RMC1 | ZBTB8OS | TMCC2 |
| 46279 | FAM13A_AS1 | PI4K2A | PPIL2 | LOC440311 | CSDC2 |
| MAP3K13 | USP16 | LAMA4 | PVR | TRMT10A | BCL3 |
| HSD11B1 | MLLT10 | PPM1H | ZNF182 | CCNI | SZT2 |
| ZNF767P | LOC151174 | KCNA3 | MROH8 | NEURL2 | HSD17B11 |
| ARHGEF10L | IER2 | CPLX2 | HSPBAP1 | RALGAPB | PIAS2 |
| MPHOSPH9 | KYAT1 | SLC39A14 | DDX55 | CCDC39 | NECAB3 |
| TRPM3 | RHOBTB2 | SLC38A1 | RCC2 | CORO6 | DSP |
| PHF8 | CEMIP | TSPAN1 | LINC02210 | PPTC7 | SCX |
| NFKBIZ | KCNT1 | TRNP1 | YBEY | STK26 | RNF168 |
| PARD6B | MYO5C | GPHN | MAML1 | NUTM2B_AS1 | HIST1H1A |
| SLC47A1 | INTS9 | SLC15A4 | SHMT1 | ST8SIA1 | SLC8A1 |
| MAGI3 | GIGYF1 | ZNF287 | TMEM14C | LRRC38 | KIAA1551 |
| HCG4 | GHRLOS | RPUSD2 | LINC00889 | RAD52 | DPY19L2 |
| IL6R | AP4E1 | KLHL24 | ZFYVE9 | TMEM265 | FKBP7 |
| REEP2 | AASDH | TAS2R31 | NDUFC1 | SYNDIG1 | NR1D1 |
| PRKCB | RRN3P3 | LOC100133091 | TSEN54 | ACTN4 | RAB40AL |
| RASIP1 | FGD5 | TMEM229B | DPY19L2P4 | DGCR9 | SLC25A34 |
| REPS1 | KANSL1 | SLC25A36 | ABCC8 | ARID4B | RNF157 |
| PACSIN3 | DCAF8L2 | ABCG4 | LSMEM1 | H1F0 | THSD4 |
| OIP5_AS1 | P3H1 | AFF1 | C3orf18 | DNAAF5 | FAM135B |
| MKLN1 | METAP2 | LOC643406 | CHD2 | JAKMIP3 | HERPUD1 |
| PIP5KL1 | SEC14L5 | EIF5 | COQ3 | PPIG | MAP3K9 |
| RGL2 | KDELC1 | ALKBH8 | GNPAT | FES | CTTNBP2 |
| MRPS5 | AKAIN1 | SEMA6D | TMEM223 | DOPEY2 | CDK14 |
| MBL1P | SF3B1 | KNG1 | ERMP1 | CASP3 | CLN6 |
| MPC1 | CA7 | BMS1 | PLEKHA6 | SYT15 | DCLRE1C |
| SLC25A17 | CDON | ZBTB7A | MROH5 | KCNS1 | KIZ |
| GNAS | CCDC138 | INSR | LOC285074 | RRM2B | PRR22 |
| HAPLN4 | PHTF1 | GABPB1_IT1 | CAMK2G | LINC00652 | PRDM11 |
| ZNF714 | DNMT3A | CLEC4M | LIN28B | NGLY1 | ANKZF1 |
| RBFOX3 | ZBTB43 | BTN2A2 | TRIM44 | TTC39A | NDUFS1 |
| SMCHD1 | MICAL3 | ARL5A | RBM7 | BAAT | MAPKBP1 |
| NARS | WDPCP | SPECC1L | COL19A1 | RYR2 | HELQ |
| CISD2 | TBC1D1 | TMEM255B | ZMYM6 | FAM78B | WASHC1 |
| CAMK1D | BACH2 | KCNB1 | RCC1L | FAM167B | WAPL |
| ITGA11 | MRNIP | ZNF407 | LTB | SLC6A6 | RP9 |
| CHGA | RFPL2 | ASAH2 | ANKRD13A | IGDCC3 | SLC20A2 |
| CALCOCO1 | SEPHS2 | CFL2 | PILRA | FKBP5 | SORCS1 |
| RAD51_AS1 | ZNF655 | DCLK1 | NOB1 | ZFAT | FAM118A |
| SMG9 | REPIN1 | CCNE1 | ZNF136 | PLEKHO1 | ZMYM5 |
| ZNF500 | MEIKIN | SPIDR | ZBTB16 | GPR108 | JARID2 |
| ZNF446 | KCNH4 | SPHK1 | RALGAPA1 | SLC33A1 | CMPK1 |
| ELMO3 | PDE7A | ARL5B | CACNG8 | UBE2O | PPP3CA |
| HMBS | HSF4 | PCDHB16 | EMX1 | NEFH | EZH2 |
| UNC13C | MCM8 | SPECC1 | MXI1 | LUZP1 | MATN4 |
| THG1L | KIF9_AS1 | UNKL | FAM78A | ZNF385B | KMT5B |
| ANKFY1 | EHMT1 | KCNK10 | CCDC84 | DTNB | PNKD |
| TRMU | SMYD4 | ZNF223 | STRADA | SDHAF2 | JAKMIP2 |
| GGT7 | RIF1 | GAPVD1 | HELB | HERC2P2 | GNPDA2 |
| LAPTM4B | TLX2 | VN1R1 | UBN2 | MYNN | CHD1L |
| MTBP | KMT2E | LOC100129917 | TCHH | ZNF138 | TTI2 |
| KRI1 | HES6 | C17orf75 | NAT8L | CLCN4 | ARMC8 |
| RSBN1L | PCNA | TOP1P2 | AMD1 | CBX7 | SMOC2 |
| ZNF785 | SYNE1 | ATP1A4 | KCTD8 | ITPR1 | CNTLN |
| MYC | DEPTOR | CLCN2 | CDKN2C | LINC01106 | RCN3 |
| ACVR1C | CCDC47 | SUPT16H | SPIN3 | MSANTD2 | CEP120 |
| FAN1 | ASTN2 | SULT1A2 | HNRNPF | ST7L | TDP2 |
| LINC02012 | PRC1 | HNRNPR | ZBTB21 | CEP97 | RASA3 |
| ARL13B | KIF5A | EPB41L5 | DOK3 | ZKSCAN4 | WASHC2C |
| ZNF563 | IL15RA | ZNF234 | MTAP | FGF7 | CNST |
| EML2 | ZNF629 | PPT2 | TMEM35A | RTEL1 | PPP1R3D |
| CHFR | OGDHL | FCHO2 | PGBD4 | CENPBD1 | TMEM120B |
| MRPL48 | ASAH2B | ZNF284 | ZNF620 | PPWD1 | MAML3 |
| FBXO21 | CUL3 | PCNX2 | FANCM | TAS2R5 | TMEM161B_AS1 |
| ZNF251 | DNASE1L2 | GSAP | PLEKHM1 | FANCC | APOL2 |
| DAZAP1 | USF2 | TAF3 | GPR158 | ZNF180 | EDNRA |
| AGAP5 | PRRT3 | MFAP3 | ALG2 | WASHC4 | SPINK8 |
| HCN2 | NFYC | DFFB | USP33 | NFATC3 | AIFM3 |
| CCDC126 | AMH | RCL1 | NUP205 | ARSG | SENP7 |
| FAM57B | GLRX | ZBED9 | ADGRL2 | CDYL | LRPPRC |
| MIR9_3HG | ARHGEF17 | FGF18 | VWDE | POGLUT1 | TNFRSF12A |
| ANKRD20A5P | PTPRD | GPR85 | ESRRB | TSC22D3 | JAK1 |
| MITF | IL17RE | ZBTB8A | ZGRF1 | SPACA6 | SLTM |
| LINC01089 | MED1 | MTFP1 | MADCAM1 | SLC22A23 | CUX2 |
| ZNF662 | AMT | PPP3CC | EPC1 | ZNF8 | NELFA |
| PUDP | ZSCAN31 | ZNF322 | NRG1 | AP1S2 | OLFML2A |
| NUDT8 | ARID5B | PDPR | ZNF782 | AGO4 | TPCN1 |
| MRPS11 | TANC1 | KHDC4 | B4GALT6 | HSD11B2 | PRH2 |
| HSPB6 | ST6GALNAC1 | SEPT7P2 | EMCN | MYO15A | EMILIN3 |
| TTC17 | MED23 | LINC01000 | FAM209A | ITPRIPL1 | IRF3 |
| ATP6V1E2 | TSHZ1 | MIER1 | ZNF354C | EMB | ZBTB34 |
| TAZ | MET | DPP8 | INA | MINDY3 | TNRC6A |
| MICA | FAM106A | SCAF11 | UVRAG | LOC728554 | MYBBP1A |
| DPM3 | PCDH11Y | DNA2 | TNNC2 | NSRP1 | THSD7A |
| RPS6KB1 | ABCA9 | PRKAB1 | SCARF2 | ZNF232 | KAT8 |
| ANKRD20A4_ANKRD20A20P | IFNAR2 | SKA2 | ABCA7 | POU6F2 | MCM7 |
| CIT | IFFO1 | EDEM1 | LYPLA1 | PIK3CA | MAP4K2 |
| CPNE9 | ANAPC4 | JRK | ERRFI1 | RBM15 | MAPK6 |
| RADIL | RING1 | CD2AP | PPM1B | MYMX | IFFO2 |
| NKX3_1 | LOC100286906 | DCLRE1A | PROSER1 | TRAK2 | TBC1D8 |
| THOC2 | NKX1_2 | ACTR1B | ZNRD1 | TUBA8 | MFAP3L |
| ZBTB40 | BICDL2 | ABCA5 | SNW1 | DPH7 | DDHD2 |
| RACGAP1 | DEXI | LINC01405 | PAN2 | INPP5A | HCN1 |
| B3GNT9 | MCPH1 | PRPF39 | MICB | PCDH12 | RBBP8 |
| SBNO1 | RNF144A | NREP | MRPS31 | ZNF562 | BCAS4 |
| RAD54L | ARNTL | ZNF493 | LOC100288203 | ORAI2 | LRRC8C |
| LIAS | FSBP | CRTC2 | CWC22 | CDKN2AIP | PLAU |
| SORT1 | ZNF365 | GOLGA8A | ZC3H18 | CRYGS | NFIX |
| TAS2R4 | MAP2K6 | GPR180 | CAPN7 | C3orf58 | NVL |
| DYNC2H1 | HRC | TAP1 | HS6ST2 | MON2 | TCF7L2 |
| ZNF35 | SAT2 | WASHC2A | CCZ1 | IQUB | PACSIN2 |
| ABCB9 | INO80 | RFX5 | ABLIM2 | TARS | HNRNPDL |
| KLHL21 | FAM225B | TAF2 | POMT2 | IRAK1BP1 | WDR4 |
| KLHL15 | RASSF8_AS1 | STK36 | C15orf41 | MAP3K14 | SLFN11 |
| MTR | MAGEL2 | PPP1CC | MGA | PIP4P2 | PCBP2 |
| DTX3 | SUMF2 | FBXL20 | CABLES2 | BLMH | FZD6 |
| SLC17A5 | CLASRP | ST3GAL5 | ATG4D | GANC | IGF1R |
| CSKMT | TAS2R20 | RAD54L2 | LONRF3 | DDX3X | PPP1R3B |
| LINC01128 | WIPI1 | ZDHHC13 | GPR137C | FBXO33 | PRMT9 |
| FLRT3 | OTUD6B_AS1 | UBE4B | CDK19 | KLC2 | LINC00667 |
| FRMD5 | SETDB1 | MRPL34 | PALB2 | ZDHHC18 | RBM48 |
| NEGR1 | DXO | SDHA | GRPEL2 | GID8 | LRRFIP1 |
| APIP | WNT3 | PAGR1 | STXBP5L | N4BP2 | CCDC144B |
| KCND3 | UBE2G1 | SEPHS1 | SLC38A5 | KCNV2 | HDAC4 |
| SUPV3L1 | PCTP | SLC46A3 | TMEM109 | GK | KLHL28 |
| SIRT7 | POLG2 | USP8 | SUN1 | GRIN2A | SEC24B |
| RRM1 | FBLN7 | CTSK | TMEM250 | TBCEL | FAP |
| SLC44A5 | ATAD2B | GDPD1 | TSPYL5 | DET1 | TAS2R43 |
| C8orf59 | ZNF133 | SLCO4A1 | UBAC2 | ATP6V0A2 | ZNF3 |
| SLC7A5P2 | MAP6D1 | TENM2 | SRRM4 | TAP2 | RTKN2 |
| VSTM2A | RAB30 | TYW1B | LRRK2 | FMN1 | ARHGAP32 |
| ISY1 | BCOR | CLN3 | TOR2A | MAX | FUT11 |
| TMEM45B | HIVEP2 | SAR1B | PLAT | TBC1D31 | XPO4 |
| NELFCD | HECTD1 | IGLON5 | ARC | NRIP2 | COL6A1 |
| H2AFX | ODF2 | LOC284009 | EP300 | HERC2P9 | DCAF1 |
| MID2 | GOLGA4 | PCGF6 | MRPS36 | KCMF1 | FIZ1 |
| TRIP4 | GREB1L | IMPA1 | EMC3_AS1 | FAM133B | IGFBP2 |
| ULK3 | FOCAD | CNNM1 | KIF2C | ST3GAL2 | NEAT1 |
| ZNF783 | GPATCH1 | SLC45A1 | AGO3 | CCAR1 | MLPH |
| AFTPH | KLHL8 | TNPO1 | LOC101926935 | TMCC1 | CCDC144CP |
| ARIH2 | HLTF | ORC4 | FAM126B | SRGAP2 | ZNF709 |
| SLC25A5 | RAB3GAP2 | EEF1AKMT2 | LINC01158 | RBMS1 | ZNF567 |
| BEND7 | ARHGAP39 | OR2L3 | CARD19 | ZSWIM5 | GFM1 |
| ELL | PAX8_AS1 | NUP98 | LGR6 | ROCK2 | ASH1L |
| PXN | SIKE1 | BCL9 | FLYWCH1 | MRPL45 | IRS1 |
| DHX33 | WDR70 | ARL17A | COL11A2 | THOC5 | KNDC1 |
| PLOD2 | ZNF598 | STAU2 | CD99L2 | KIN | BRD2 |
| LOXL2 | ZBTB5 | MTHFD2L | NEFM | CALML4 | SH3BGRL |
| NXF1 | C1orf162 | E2F6 | GALE | SEC62 | ATG4B |
| SLC26A10 | PPARGC1B | NAXD | SERTAD3 | LRIG2 | NDUFA10 |
| SLC4A8 | SAA2 | LYG1 | ADAMTSL1 | ZNF487 | C2orf68 |
| ZNF30 | ANKRD49 | IL1RAP | ZBTB47 | RABL6 | CHD6 |
| GLA | ABL2 | RALB | MFF | XIRP1 | INTS6L |
| RNFT1 | ULK1 | FLCN | DEPDC1 | MTG2 | HNRNPA3P1 |
| TRIM37 | SDSL | ZNF711 | UBE2G2 | TMEM132C | TRIM35 |
| JMY | SFI1 | CHCHD3 | PDSS1 | OXNAD1 | ACYP2 |
| RAD51C | WDR91 | GCNA | GAS6 | KCNJ11 | ALKBH1 |
| MTO1 | PCDHGB4 | GNL1 | CBY1 | HYDIN2 | FAM84A |
| ZFX | MAP2K3 | IFIH1 | GLE1 | ADAM9 | MDC1 |
| USPL1 | PLK1 | AGAP11 | UTP15 | SPRY3 | USP31 |
| PFKFB2 | P2RX6 | DUBR | ALKBH5 | ELP6 | ZCCHC9 |
| ZFP37 | EPC2 | DIP2A | SLC35G5 | BRINP2 | TSTD2 |
| FAM133CP | ABHD6 | DSTYK | ZNF669 | LARP4 | ATP6AP1L |
| PDIK1L | MPP7 | RAB40A | SORL1 | FAM208B | CNEP1R1 |
| ZNF491 | ATP1B3 | PEG3 | RPL15 | TAS2R45 | AKAP17A |
| SAV1 | ZNF781 | SETBP1 | FNDC4 | POLK | PAQR3 |
| COIL | SPEN | ESM1 | ZBED1 | RNF115 | ARHGAP27 |
| UBE2CP5 | ZC3H10 | BARD1 | CAMK2N2 | ZNF92 | IER5 |
| MAPK10 | ZNF44 | CCDC134 | KDM2B | RGS7 | TADA2A |
| ANAPC10 | ASPRV1 | METTL17 | NUDT6 | NPIPB11 | ZNF417 |
| FMN2 | KCNN4 | AMY2A | LIN7A | SETD2 | TOP3A |
| RABEP2 | PTPN4 | SPOCD1 | TRMT11 | RWDD2B | RBFOX1 |
| UBAP1 | KIF5B | SMIM13 | RTN4 | LINC00115 | LINC02145 |
| KAZALD1 | AGPAT3 | CRTC3 | SRXN1 | ATP11B | ZNF343 |
| BUB3 | TOP3B | HOXD9 | ALS2 | CBWD5 | NLRP3 |
| NUP210 | LGI3 | PLAUR | OXA1L | KSR2 | CDC42SE1 |
| MCU | C2orf54 | REEP6 | SNHG17 | PMEPA1 | TARBP2 |
| RABGEF1 | RBM39 | CACNA1C | GPR135 | B4GALT4 | SPOPL |
| SMARCC1 | CHML | TAS2R19 | EMBP1 | CAB39 | KRTAP5_7 |
| EPB41L1 | NPHP4 | LPGAT1 | MRPL22 | ABHD17A | SYBU |
| TSNARE1 | LINC01596 | LEMD3 | MTFR1 | CAMKK1 | NXPE3 |
| REL | CLOCK | ATP11A | HOMER3 | CYTH1 | RAI1 |
| ZNF879 | CCND2_AS1 | HYDIN | NIPA1 | GOLGA6L10 | ABCB7 |
| NAGPA | MAP3K10 | TRPM7 | CEP250 | OR2AK2 | LOC100505938 |
| BHLHE22 | GOLGA2P5 | SESN2 | MDN1 | NUMA1 | ZKSCAN7 |
| NUDT12 | CD209 | ZNF791 | SEMA5B | TTLL11 | ATXN2L |
| ARRDC1_AS1 | OTUD7A | FGF7P3 | USP6NL | GABRB2 | ADGRL4 |
| RPAP2 | C8orf31 | ABHD12 | CTCF | LINC00266_1 | LOC105374727 |
| LOC100190986 | GSE1 | TET1 | OR2L13 | MAT2B | LOC100287497 |
| C9orf72 | LOC283788 | BRICD5 | LRRC75A | CACNG7 | FBXO42 |
| LMO2 | SPDYE5 | CCNL1 | LARP6 | SLC25A27 | ATXN7L2 |
| HIST4H4 | BMS1P4 | ANKRD18CP | SENP1 | NPR3 | HDAC8 |
| DNAJC28 | USP3 | BID | PPP2R5D | SLC25A15 | TMEM39A |
| TRRAP | UPF1 | RABL2A | KLHL11 | XBP1 | RAB11B |
| CELSR2 | ASPSCR1 | TMEM249 | FAM193A | TMF1 | GUSBP3 |
| PWAR5 | ALDH18A1 | PRAG1 | TGFBR1 | GTF3A | EPHB2 |
| DPT | KIAA0586 | WDFY3 | PIDD1 |  |  |

**Table S4.5** The results of Gene Ontology (GO) enrichment analysis

| **Abbreviation  (in Fig 4d)** | **Full GO Term** | **GO ID** | ***p*_FDR_** | **gene counts** | **Gene rasio** |
| --- | --- | --- | --- | --- | --- |
| MF | Molecular Function |  |  |  |  |
| BP | Biological Process |  |  |  |  |
| CC | Cellular Component |  |  |  |  |
| downregulated genes from ALE & ALE-FCNM | | | | | |
| MF: steroid binding | MF: steroid binding | GO:0005496 | **0.021** | 12 | 0.09 |
| MF: ICTTA | MF: inorganic cation transmembrane transporter activity | GO:0022890 | **0.021** | 35 | 0.05 |
| MF: EREB | MF: estrogen response element binding | GO:0034056 | **0.021** | 4 | 0.36 |
| MF: ECTTA | MF: monoatomic cation transmembrane transporter activity | GO:0008324 | **0.021** | 36 | 0.05 |
| MF: MITTA | MF: metal ion transmembrane transporter activity | GO:0046873 | **0.003** | 33 | 0.06 |
| CC: calyx of Held | CC: calyx of Held | GO:0044305 | **<0.001** | 8 | 0.22 |
| upregulated genes from ALE & ALE-FCNM | | | | | |
| MF: CIB | MF: cadmium ion binding | GO:0046870 | **<0.001** | 6 | 0.50 |
| MF: MICA | MF: monoatomic ion channel activity | GO:0005216 | **<0.001** | 49 | 0.08 |
| MF: PPTTA | MF: passive transmembrane transporter activity | GO:0022803 | **<0.001** | 52 | 0.08 |
| MF: channel activity | MF: channel activity | GO:0015267 | **<0.001** | 52 | 0.08 |
| MF: CIO | MF: copper ion binding | GO:0005507 | **<0.001** | 14 | 0.18 |
| BP: SO | BP: synapse organization | GO:0050808 | **<0.001** | 60 | 0.08 |
| BP: CRTMI | BP: cellular response to zinc ion | GO:0071294 | **<0.001** | 11 | 0.38 |
| BP: SRTMI | BP: stress response to metal ion | GO:0097501 | **<0.001** | 10 | 0.45 |
| BP: DOIC | BP: detoxification of inorganic compound | GO:0061687 | **<0.001** | 11 | 0.44 |
| BP: CCS | BP: cell-cell signaling | GO:0007267 | **<0.001** | 112 | 0.07 |
| CC: PM | CC: postsynaptic membrane | GO:0045211 | **<0.001** | 44 | 0.11 |
| CC: SM | CC: synaptic membrane | GO:0097060 | **<0.001** | 59 | 0.10 |
| CC: postsynapse | CC: postsynapse | GO:0098794 | **<0.001** | 86 | 0.08 |
| CC: NP | CC: neuron projection | GO:0043005 | **<0.001** | 121 | 0.07 |
| CC: presynapse | CC: presynapse | GO:0098793 | **<0.001** | 83 | 0.09 |
| downregulated genes from SDM-FCNM | | | | | |
| MF: PMAA | MF: protein-macromolecule adaptor activity | GO:0030674 | **<0.001** | 177 | 0.13 |
| MF: MAA | MF: molecular adaptor activity | GO:0060090 | **<0.001** | 190 | 0.12 |
| MF: DNA-HA | MF: DNA helicase activity | GO:0003678 | **<0.001** | 21 | 0.27 |
| MF: CA-DNA | MF: catalytic activity, acting on DNA | GO:0140097 | **<0.001** | 48 | 0.18 |
| MF: HA-AA | MF: hydrolase activity, acting on acid anhydrides | GO:0016817 | **<0.001** | 158 | 0.12 |
| BP: DNA-DR | BP: DNA damage response | GO:0006974 | **0.004** | 124 | 0.13 |
| BP: mRNA processing | BP: mRNA processing | GO:0006397 | **0.004** | 79 | 0.14 |
| BP: DNA repair | BP: DNA repair | GO:0006281 | **0.007** | 88 | 0.13 |
| BP: RNA splicing | BP: RNA splicing | GO:0008380 | **0.007** | 72 | 0.14 |
| BP: DNA-MP | BP: DNA metabolic process | GO:0006259 | **0.011** | 131 | 0.12 |
| CC: nuclear body | CC: nuclear body | GO:0016604 | **<0.001** | 132 | 0.14 |
| CC: nuclear speck | CC: nuclear speck | GO:0016607 | **<0.001** | 67 | 0.15 |
| CC: NRG | CC: nuclear ribonucleoprotein granule | GO:0140168 | **0.002** | 73 | 0.14 |
| CC: RG | CC: ribonucleoprotein granule | GO:0035770 | **0.002** | 99 | 0.13 |
| CC: catalytic complex | CC: catalytic complex | GO:1902494 | **0.003** | 204 | 0.11 |
| upregulated genes from SDM-FCNM | | | | | |
| MF: MICA | MF: monoatomic ion channel activity | GO:0005216 | **0.003** | 89 | 0.15 |
| MF: TRA | MF: transporter regulator activity | GO:0141108 | **0.003** | 41 | 0.19 |
| MF: SRB | MF: signaling receptor binding | GO:0005102 | **0.003** | 217 | 0.12 |
| MF: PDZ-DB | MF: PDZ domain binding | GO:0030165 | **0.003** | 28 | 0.23 |
| MF: OA | MF: oxidoreductase activity | GO:0016491 | **0.003** | 105 | 0.14 |
| BP: ATS | BP: anterograde trans-synaptic signaling | GO:0098916 | **<0.001** | 164 | 0.17 |
| BP: TS | BP: trans-synaptic signaling | GO:0099537 | **<0.001** | 167 | 0.17 |
| BP: SO | BP: synapse organization | GO:0050808 | **<0.001** | 138 | 0.19 |
| BP: synaptic signaling | BP: synaptic signaling | GO:0099536 | **<0.001** | 175 | 0.18 |
| BP: cell-cell signaling | BP: cell-cell signaling | GO:0007267 | **<0.001** | 261 | 0.16 |
| CC: glutamatergic synapse | CC: glutamatergic synapse | GO:0098978 | **<0.001** | 162 | 0.19 |
| CC: synaptic membrane | CC: synaptic membrane | GO:0097060 | **<0.001** | 129 | 0.21 |
| CC: postsynapse | CC: postsynapse | GO:0098794 | **<0.001** | 196 | 0.18 |
| CC: neuron projection | CC: neuron projection | GO:0043005 | **<0.001** | 274 | 0.16 |
| CC: presynapse | CC: presynapse | GO:0098793 | **<0.001** | 176 | 0.19 |

**Appendix 5 Receptor association analysis**

**Table S5.1** Receptor association analysis results

| Receptor | ALE | | ALE-FCNM | | SDM-FCNM | | SDM | |
| --- | --- | --- | --- | --- | --- | --- | --- | --- |
|  | *r* | *p_Bonferroni_* | *r* | *p_Bonferroni_* | *r* | *p_Bonferroni_* | *r* | *p_Bonferroni_* |
| 5HT1a | 0.23 | **0.002** | 0.30 | **<0.001** | 0.28 | **<0.001** | -0.03 | 1.000 |
| 5HT1b | 0.11 | 1.000 | 0.09 | 1.000 | 0.24 | **<0.001** | -0.07 | 1.000 |
| 5HT2a | 0.22 | **0.002** | 0.21 | **0.004** | 0.23 | **<0.001** | -0.06 | 1.000 |
| 5HT4 | 0.24 | **<0.001** | 0.22 | **0.002** | 0.23 | **0.001** | -0.01 | 1.000 |
| 5HT6 | 0.13 | 0.386 | 0.19 | **0.015** | 0.17 | 0.053 | -0.08 | 1.000 |
| 5HTT | 0.12 | 0.783 | 0.17 | **0.048** | 0.03 | 1.000 | -0.04 | 1.000 |
| A4B2 | 0.04 | 1.000 | 0.15 | 0.217 | 0.30 | **<0.001** | -0.04 | 1.000 |
| CB1 | 0.20 | **0.008** | 0.27 | **<0.001** | 0.40 | **<0.001** | -0.09 | 1.000 |
| D1 | 0.26 | **<0.001** | 0.45 | **<0.001** | 0.35 | **<0.001** | -0.02 | 1.000 |
| D2 | 0.16 | 0.086 | 0.13 | 0.545 | 0.14 | 0.361 | 0.05 | 1.000 |
| DAT | 0.19 | **0.019** | 0.29 | **<0.001** | 0.18 | **0.031** | 0.04 | 1.000 |
| GABAa | -0.05 | 1.000 | -0.06 | 1.000 | -0.13 | 0.465 | -0.05 | 1.000 |
| H3 | 0.25 | **<0.001** | 0.43 | **<0.001** | 0.53 | **<0.001** | -0.01 | 1.000 |
| M1 | 0.10 | 1.000 | 0.08 | 1.000 | 0.13 | 0.378 | -0.07 | 1.000 |
| mGluR5 | 0.23 | **0.001** | 0.33 | **<0.001** | 0.38 | **<0.001** | -0.03 | 1.000 |
| MOR | 0.33 | **<0.001** | 0.45 | **<0.001** | 0.63 | **<0.001** | 0.04 | 1.000 |
| NET | -0.18 | **0.025** | -0.25 | **<0.001** | -0.21 | **0.006** | -0.05 | 1.000 |
| NMDA | 0.14 | 0.313 | 0.20 | **0.008** | 0.09 | 1.000 | 0.02 | 1.000 |
| VAChT | 0.05 | 1.000 | 0.17 | 0.053 | 0.17 | 0.063 | -0.03 | 1.000 |

**Appendix 6 Robustness analysis of multiple linear regression**

**Table S6.1** Robustness analysis of multiple linear regression

|  | Independent  variable | dependent variable | *β* | OLS *p* | Bonferroni *p* | FDR *p* | HC3 *p* | Bootstrap *p* | Perm *p* | Outlier *p* |
| --- | --- | --- | --- | --- | --- | --- | --- | --- | --- | --- |
| Gradient 1 | FCG | ALE | 0.546 | **<0.001** | **<0.001** | **<0.001** | **<0.001** | **<0.001** | **0.001** | **<0.001** |
|  |  | SDM | 0.627 | **<0.001** | **<0.001** | **<0.001** | **<0.001** | **<0.001** | **0.001** | **<0.001** |
|  | MSG | ALE | -0.042 | 0.289 | 1 | 0.289 | 0.276 | 0.298 | 0.541 | 0.237 |
|  |  | SDM | 0.138 | **<0.001** | **<0.001** | **<0.001** | **<0.001** | **<0.001** | **0.046** | **<0.001** |
|  | TSG | ALE | 0.175 | **0.015** | 0.062 | **0.021** | 0.064 | 0.06 | 0.172 | 0.121 |
|  |  | SDM | 0.111 | 0.062 | 0.248 | 0.083 | 0.1 | 0.082 | 0.381 | 0.054 |
|  | RSG | ALE | 0.187 | **0.009** | **0.037** | **0.018** | **0.031** | **0.014** | 0.158 | **0.005** |
|  |  | SDM | 0.1 | 0.092 | 0.366 | 0.092 | 0.125 | 0.102 | 0.433 | 0.194 |
| Gradient 2 | FCG | ALE | 0.721 | **<0.001** | **<0.001** | **<0.001** | **<0.001** | **<0.001** | **0.001** | **<0.001** |
|  |  | SDM | 0.503 | **<0.001** | **<0.001** | **<0.001** | **<0.001** | **<0.001** | **0.001** | **<0.001** |
|  | MSG | ALE | -0.032 | 0.457 | 1 | 0.457 | 0.446 | 0.498 | 0.645 | 0.148 |
|  |  | SDM | 0.13 | **0.006** | **0.024** | **0.012** | **0.006** | **0.006** | **0.045** | **0.026** |
|  | TSG | ALE | 0.177 | **0.02** | 0.081 | **0.04** | **0.013** | **0.006** | 0.142 | **0.015** |
|  |  | SDM | 0.213 | **0.011** | **0.044** | **0.015** | **0.008** | **0.01** | 0.07 | **<0.001** |
|  | RSG | ALE | -0.107 | 0.172 | 0.688 | 0.229 | 0.177 | 0.166 | 0.379 | 0.45 |
|  |  | SDM | -0.056 | 0.514 | 1 | 0.514 | 0.477 | 0.45 | 0.65 | 0.295 |
| Gradient 3 | FCG | ALE | 0.091 | 0.211 | 0.844 | 0.211 | 0.235 | 0.22 | 0.335 | 0.052 |
|  |  | SDM | 0.132 | 0.103 | 0.411 | 0.206 | 0.115 | 0.136 | 0.15 | **0.017** |
|  | MSG | ALE | 0.175 | **<0.001** | **<0.001** | **<0.001** | **<0.001** | **<0.001** | **0.009** | **<0.001** |
|  |  | SDM | -0.019 | 0.737 | 1 | 0.737 | 0.715 | 0.688 | 0.772 | 0.316 |
|  | TSG | ALE | 0.113 | 0.204 | 0.818 | 0.211 | 0.228 | 0.234 | 0.332 | 0.272 |
|  |  | SDM | 0.123 | 0.211 | 0.845 | 0.282 | 0.213 | 0.22 | 0.265 | 0.487 |
|  | RSG | ALE | 0.372 | **<0.001** | **<0.001** | **<0.001** | **<0.001** | **<0.001** | **0.005** | **<0.001** |
|  |  | SDM | 0.305 | **0.005** | **0.021** | **0.021** | **0.007** | **0.004** | **0.018** | **0.001** |

Note. The dependent variables “ALE” and “SDM” refer to the ALE-FCNM-based functional connectivity gradient and the SDM-FCNM-based functional connectivity gradient, respectively. FCG, functional connectivity gradient; MSG, morphometric similarity gradient; TSG, transcriptomic similarity gradient; RSG, receptor similarity gradient. Bold text indicates *p* < 0.05
